# PAXIP1-PAGR1 directs cohesin recruitment during break-induced telomere repair

**DOI:** 10.64898/2026.09.14.751543

**Authors:** Soo-Yeon Hwang, Xiaoying Wu, Xiangao Huang, Fei Li, Zihua Wang, Eun Young Yu, Yunxia Xu, Zhengyu Zhang, Inhye Moon, Seul-Ah Kim, Boris Yamrom, Sieun Yang, Yizhe Wang, Youngjoo Kwon, Neal F. Lue, Jihye Paik, Changjiang Dong, Hongwu Zheng

**Affiliations:** Department of Pathology and Laboratory Medicine, Weill Cornell Medicine, New York, NY 10065, USA; Key Laboratory of Combinatorial Biosynthesis and Drug Discovery, Ministry of Education, School of Pharmaceutical Sciences, Wuhan University, Wuhan 430071, China; Cold Spring Harbor Laboratory, Cold Spring Harbor, NY 11724, USA; Department of Neurosurgery, Southwest Hospital, Chongqing, 400038, China; Department of Microbiology and Immunology, W. R. Hearst Microbiology Research Center, Weill Cornell Medicine, New York, NY 10065, USA; College of Pharmacy and Graduate School of Pharmaceutical Sciences, Ewha Womans University, Seoul 03760, Republic of Korea; Graduate Program in Innovative Biomaterials Convergence, Ewha Womans University, Seoul 03760, Republic of Korea; Department of Cardiovascular Medicine, The Second Affiliated Hospital of Wenzhou Medical University, Wenzhou, 325024, China

**Author notes:** Correspondence (J.P.), (C.D.), & (H.Z.). These authors contributed equally to this work (Soo-Yeon Hwang, Xiaoying Wu, Xiangao Huang).

## Abstract

Cohesin is a conserved multiprotein complex (SMC1, SMC3, RAD21, and either STAG1 or STAG2) that organizes three-dimensional genome architecture and regulates chromosome segregation, gene expression, and DNA damage repair^1–4^. Following double-strand breaks (DSBs), cohesin is recruited to sites of DNA damage - a process considered essential for efficient homologous recombination^5–15^. Yet how DSB signaling elicits cohesin recruitment and subsequent cohesion establishment remains poorly understood. Here we show that telomere replication stress activates *de novo* STAG2-cohesin loading, thereby promoting break-induced telomeric DNA repair. We demonstrate that this DNA break-elicited cohesin recruitment is strictly controlled by the BRCT domain-containing DNA damage recognition factor PAXIP1 and its functional partner PAGR1. Cryo-electron microscopy structure reveals that PAGR1, together with PAXIP1, physically binds to a composite interface formed by the STAG2-RAD21 cohesin subcomplex. Complementary mutational and biochemical analyses define the molecular basis of this interaction and establish its essential role in break-induced cohesion establishment. Furthermore, we show that PAXIP1-PAGR1-enacted STAG2-cohesin recruitment complements with the PML body-associated pathway in orchestrating break-induced alternative lengthening of telomeres (ALT). Concurrent depletion of PML together with PAXIP1, PAGR1 or STAG2 disrupts ALT-mediated telomere maintenance, leading to end-to-end chromosomal fusion and mitotic cell death. Collectively, these findings uncover a distinctive molecular mechanism through which DSB signaling directs *de novo* cohesion establishment, and highlight its critical importance in break-induced telomere repair.

## Introduction

The repair of chromosomal double-strand breaks (DSBs) via homologous recombination (HR) is essential for maintaining genomic stability ^16, 17^. Among HR pathways, break-induced replication (BIR) repairs one-ended DSBs through long-tract, homology-directed DNA synthesis. BIR underpins HR-directed alternative lengthening of telomeres (ALT) ^18–22^, a telomerase-independent telomere length maintenance mechanism that supports ∼10%-15% of human cancers. Unlike HR-mediated repair of two-ended DSBs, which preferentially uses tethered sister chromatids as repair templates ^23, 24^, ALT-associated telomere repair by BIR is thought to initiate through strand invasion, often into a non-sister telomere duplex, followed by DNA synthesis extending to the chromosome end ^19, 22, 25^. For such long-tract DNA repair synthesis, the broken termini must be brought into proximity with a homologous template and maintained in physical association throughout repair. Accordingly, a subset of ALT telomeres coalesces broken DNA termini, templates, and repair factors into the characteristic ALT-associated PML nuclear bodies (APBs), which provide a recombinogenic microenvironment assisting telomere repair ^26–29^. However, some ALT-positive cells maintain their telomere length independently of APB ^30, 31^, suggesting the existence of parallel mechanism(s) that guide ALT-mediated telomere repair independently of PML.

Cohesin, a ring-shaped member of the structural maintenance of chromosomes (SMC) protein family, organizes chromosomes by tethering sister chromatids and by extruding DNA loops to form topologically associating domains (TADs) ^1–3^. Cohesion is required for efficient DSB repair across species, from yeast to humans ^2, 32^. Upon DSB induction, cohesin is rapidly recruited to regions surrounding the DNA break, where it establishes targeted cohesion and localized chromatin looping ^6–9, 11, 12, 33^. Targeted cohesion promotes HR by holding a homologous donor DNA template - often the sister chromatid - in close proximity to the damaged locus, thereby facilitating homology search and preventing promiscuous repair events with distant genomic loci ^7, 8, 11–13, 34^. Meanwhile, break-anchored chromatin loops help establish DSB-induced chromatin modifications ^33^ and to participate in local homology search ^7, 12, 15^. Despite these advances, fundamental questions regarding how DSB signaling elicits cohesin recruitment remains unresolved. It is also unclear whether the DSB-induced cohesion is established through *de novo* cohesin loading or via repositioning of pre-existing cohesin complexes.

To address these questions, we combined complementary approaches to investigate BIR-mediated telomere synthesis. Our results identify the heterodimeric PAXIP1-PAGR1 complex as a critical DNA damage-sensing module that directs DSB-induced *de novo* cohesion establishment in guiding HR-mediated telomere repair.

## Results

### PML is dispensable for ALT-directed telomere DNA synthesis

ALT is characterized by the formation of multitelomere clusters and associated PML nuclear bodies ^26–29^. To define the role of PML in ALT-mediated telomere maintenance, we used CRISPR/Cas9-based genome editing to generate clonal *PML*-deletion (dPML) derivatives of U2OS cells, a widely used ALT-positive osteosarcoma cell line (Fig. 1a). Consistent with previous reports ^31, 35^, while loss of PML had little effect on U2OS cell proliferation (Extended Data Fig. 1a), its depletion significantly impaired ALT-associated telomere synthesis, as indicated by reduced 5-ethynyl-2-deoxyuridine (EdU) colocalization with telomeres in G2/M-synchronized, non-S-phase U2OS-dPML cells, relative to parental controls (Fig. 1b,c). Telomere restriction fragment (TRF) analysis of U2OS-dPML cells further revealed progressive telomere shortening during extended culture spanning 250 population doublings (PDs) (Fig. 1d). The reduction of ALT-associated telomere synthesis was specifically attributable to PML loss, as reconstitution of cDNA encoding a CRISPR-resistant and Flag-tagged PML (Flag-PMLr) restored non-S-phase telomere EdU incorporation in U2OS-dPML cells (Extended Data Fig. 1b-d), indicating that PML is required for ALT-mediated telomere maintenance in U2OS cells.

**Fig. 1.**
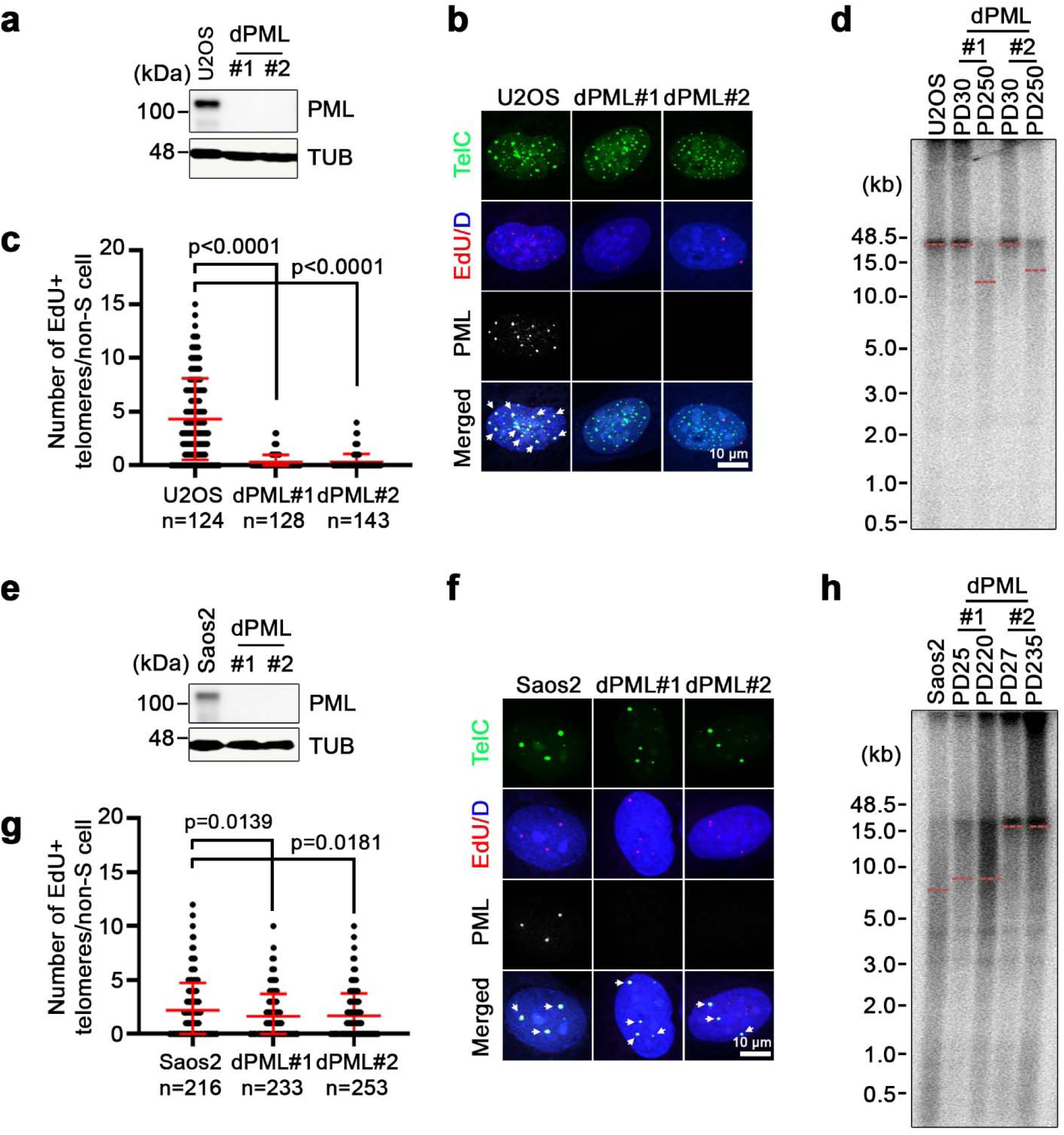
PML is non-essential for ALT telomere maintenance. **a,** Western blot analysis of PML protein expression from parental U2OS cells and U2OS CRISPR/Cas9-derived PML deletion clones (dPML#1 and #2). TUBLIN (TUB) is used as a loading control. **b,** Representative immunofluorescence (IF)-FISH images showing non-S-phase EdU colocalization with telomeres (TelG) in parental U2OS cells and U2OS-dPML clones (#1 and #2). Arrows indicate EdU-positive telomere foci. **c,** Quantification of (**b**), showing the number of non-S-phase EdU-positive telomere foci per cell in parental U2OS cells and dPML clones. Data represent the mean ± s.e.m. from three independent experiments. Statistical analysis was performed using an unpaired two-tailed Student’s t test; P values are shown. **d,** Telomere restriction fragment (TRF) analysis of telomere length in parental U2OS cells and dPML clones (#1 and #2) at the indicated population doublings (PD). Genomic DNA from the indicated cells were assayed with a ^32^P-labeled TelG probe. **e,** Western blot analysis of PML protein expression from parental Saos2 cells and two PML deleted clones (dPML#1 and #2). **f,** Representative IF-FISH images showing non-S-phase EdU colocalization with telomeres (TelG) in parental Saos2 cells and dPML clones (#1 and #2). **g,** Quantification of (**f**), showing the number of non-S-phase EdU-positive telomere foci per cell in parental Saos2 and dPML clones. Data represent mean ± s.e.m. from three independent experiments. Statistical analysis was performed using an unpaired two-tailed Student’s t test; P values are shown. **h,** TRF analysis of telomere length in Saos2 parental cells and Saos2-dPML clones (#1 and #2) at the indicated population doublings (PD).

To determine whether this PML-dependency is shared by other ALT-positive cells, we next generated clonal dPML derivatives of Saos2 cells, another ALT-positive osteosarcoma cell line (Fig. 1e). In contrast to U2OS-dPML cells that exhibited progressive telomere shortening over passaging, knockout of *PML* in Saos2 cells led to only a modest reduction in non-S-phase telomere EdU incorporation relative to parental controls (Fig. 1f,g). Moreover, TRF analysis of continuously passaged Saos2-dPML cells revealed no detectable telomere attrition even after 235 PDs in culture (Fig. 1h), suggesting that PML is nonessential for telomere maintenance in Saos2 cells.

We previously developed ALT-positive cell lines from primary human lung fibroblast IMR90 following depletion of the chromatin remodeler ATRX ^36^. To assess whether PML is similarly dispensable for telomere length maintenance in these non-tumorigenic, experimentally immortalized ALT-positive cells, we generated clonal dPML derivatives of IMR-ALT#1 (Extended Data Fig. 1e). Similar to Saos2 cells, *PML* deletion in IMR90-ALT#1 cells neither provoked progressive telomere shortening nor abolished ALT-mediated telomere synthesis (Extended Data Fig. 1f-h). Together, these data suggest the existence of alternative pathway(s) that support ALT-directed telomere synthesis independently of PML.

### PAXIP1-PAGR1 promotes PML-independent ALT telomere maintenance

The finding that PML is uniquely required for telomere length maintenance in U2OS cells, but dispensable in other ALT-positive cells, suggested that U2OS might be compromised of an alternative ALT-supporting pathway. To explore this potential deficiency, we surveyed mutation profiles of ALT cell lines using the DepMap portal (https://depmap.org). This analysis identified *PAXIP1*, which encodes a BRCT domain-containing protein, as specifically mutated in U2OS cells. Sanger sequencing of PCR-amplified genomic DNA confirmed a homozygous deletion (c.2204_2214) within the *PAXIP1* coding sequence (Fig. 2a). This deletion causes a frameshift that introduces a premature stop codon 11 amino acids downstream, producing a truncated PAXIP1 protein. Copy number variations (CNV) analysis of U2OS cells using SMASH (Short Multiply Aggregated Sequence Homologies) ^37^ further revealed a hemizygous deletion encompassing chromosome 7q35-q36.3, the region containing *PAXIP1* (Fig. 2b), indicating loss of heterozygosity (LOH). Accordingly, western blot analysis detected no full-length PAXIP1 protein in U2OS cells, whereas robust expression was observed in both Saos2 and IMR90-ALT#1 cells (Fig. 2c). Notably, the absence of wild-type PAXIP1 protein in U2OS cells coincides with the lower abundance of its obligate binding partner PAGR1 protein compared to Saos2 and IMR90-ALT#1 cells, consistent with the previous report ^38^.

**Fig. 2.**
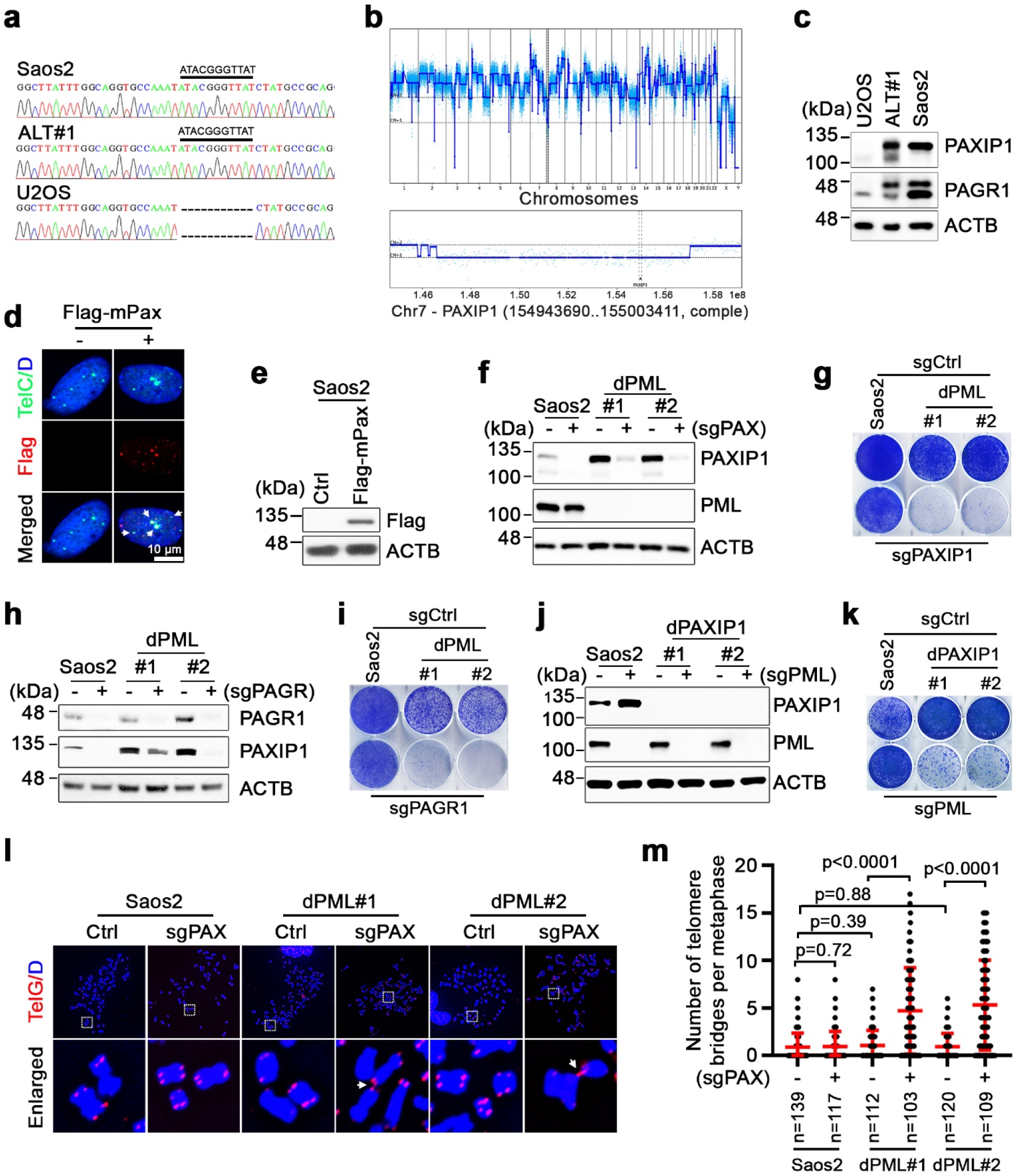
The PAXIP1-PAGR1 complex promotes PML-independent telomere maintenance in ALT cells. **a,** Sanger sequencing traces showing the *PAXIP1* deletion (c.2204_2214) in U2OS cells, compared to the wild type *PAXIP1* sequence in Saos2 and IMR90-ALT#1 cells. The indel is indicated. **b,** SMASH copy number profiles for U2OS cells. Upper panel: whole-genome view; Lower panel: expanded view of Chromosome 7, highlighting the *PAXIP1* locus (154,943,690 – 155,003,411, complement strand). **c,** Western blot analysis of PAXIP1 and PAGR1 protein expression in U2OS, IMR90-ALT#1 (ALT#1), and Saos2 cells. β-ACTIN (ACTB) serves as a loading control. **d,** Representative IF-FISH images showing colocalization of ectopically expressed, Flag-tagged murine Paxip1 (Flag-mPax, detected with anti-Flag antibody) with telomeres (TelC) in Saos2 cells. Empty vector (−) serves as a staining control. **e,** Western blot analysis confirming Flag-mPax expression in cells from (**d**). **f,** Western blot analysis of PAXIP1 and PML protein expression in parental Saos2 cells and Saos2-dPML clones (#1 and #2) following transduction with sgCtrl (−) or sgPAXIP1 (sgPAX). **g,** Crystal violet-based clonogenic assay of parental Saos2 cells and dPML clones (#1 and #2) transduced with sgCtrl or sgPAX. Crystal violet staining was performed on day 18 post-seeding. **h,** Western blot analysis of PAGR1 and PAXIP1 protein expression in parental Saos2 cells and dPML clones following transduction with sgCtrl (−) or sgPAGR1 (sgPAGR). **i,** Crystal violet-based clonogenic assay of parental Saos2 cells and dPML clones transduced with sgCtrl or sgPAGR1. Crystal violet staining was conducted on day 16 post-seeding. **j,** Western blot analysis of PAXIP1 and PML protein expression in parental Saos2 cells and CRISPR-derived PAXIP1 deletion clones (dPAX#1 and #2) following transduction of sgCtrl (−) or sgPML (+). **k,** Crystal violet-based clonogenic assay of parental Saos2 cells and dPAX clones (#1 and #2) transduced with sgCtrl or sgPML. Crystal violet staining was conducted on day 16 post-seeding. **l,** Representative telomere FISH images of metaphase spreads in parental Saos2 cells and dPML clones transduced with sgCtrl or sgPAX. Inter-chromosomal telomere fusion events are highlighted by arrows in the enlarged images. **m,** Quantification of (**l**), showing the number of telomere fusion events per metaphase in parental Saos2 cells and dPML clones transduced with sgCtrl or sgPAX. Data represent mean ± s.e.m. from three independent metaphase spread experiments. Statistical analysis was performed using an unpaired two-tailed Student’s t test. P values are shown.

PAXIP1 (also known as PAX-interacting protein 1) is a multi-BRCT domain protein implicated in DSB repair ^39–42^. Immuno-FISH analysis of Saos2 cells expressing Flag-tagged murine Paxip1 (Flag-mPax) revealed specific Paxip1 recruitment to clustered ALT telomere foci (Fig. 2d,e). To determine whether PAXIP1 mediates PML-independent ALT pathway, we depleted PAXIP1 using CRISPR/Cas9 in parental Saos2 control and *PML*-knockout Saos2-dPML cells (Fig. 2f). While PAXIP1 loss had little effect on the growth of parental Saos2 cells, its depletion in Saos2-dPML#1 and dPML#2 cells markedly suppressed their colony formation (Fig. 2g). The on-target effect of sgPAXIP1 was validated by complementing Saos2-dPML cells with sgRNA-resistant Flag-mPaxip1-encoding cDNA, which rescued the growth inhibition in sgPAXIP1-transduced Saos2-dPML cells (Extended Data Fig. 2a,b). Moreover, depletion of PAGR1 - the primary functional partner of PAXIP1 - phenocopied the effects of PAXIP1 loss on clonogenic growth in both control and Saos2-dPML cells (Fig. 2h,i). Reciprocally, we tested whether depletion of PML affected the PAXIP1-knockout cells. While PML was dispensable in parental Saos2 cells, its depletion in *PAXIP1*-knockout Saos2-dPAX cells markedly impaired their growth capacities (Fig. 2j,k). These findings indicate that PAXIP1-PAGR1 and PML pathways function independently but compensate for one another to support ALT cell growth.

This apparent synthetic lethal phenotype between PAXIP1-PAGR1 and PML promoted us to ask whether this interaction extends to non-ALT cells. To test this possibility, we generated clonal dPML derivatives from the telomerase-positive cervical cancer cell line HeLa (Extended Data Fig. 2c). Clonogenic assays showed that PML was largely dispensable for HeLa cell growth, as expected (Extended Data Fig. 2d). But unlike in Saos2-dPML cells where concomitant PAXIP1 and PML loss results in synthetic lethality, depletion of PAXIP1 in HeLa-dPML cells produced no detectable growth inhibition. Similar results were observed in telomerase-positive glioblastoma U87 cells, in which *PML* and *PAXIP1* deletions, either individually or in combination, failed to affect clonogenic growth capacities (Extended Data Fig. 2e,f), indicating that PAXIP1-PAGR1 and PML are non-essential for telomere maintenance of ALT-negative cells.

PML protein is a core structural component of APBs that facilitate ALT-mediated telomere repair ^26, 28, 29^. Given that the synthetic lethal interaction between PAXIP1-PAGR1 and PML occurs specifically in ALT-positive cells, we reasoned that these two pathways might complement one another to promote ALT-mediated telomere maintenance. Indeed, telomere FISH analysis of metaphase spreads from PAXIP1-depleted Saos2-dPML cells revealed substantially increased formation of inter-chromosomal telomere bridges, including complex fusions involving more than two chromosomal ends, as compared to their respective controls (Fig. 2l,m). By contrast, depletion of either PML or PAXIP1 individually in parental Saos2 cells did not alter metaphase telomere architecture, confirming their functional redundancy and collective importance.

We next explored whether PAXIP1 depletion affects ALT-mediated telomeric DNA synthesis. In line with PAXIP1 being dispensable for clonogenic growth of parental Saos2 cells, CRISPR/Cas9-mediated PAXIP1 depletion alone had little effect on non-S-phase EdU telomere incorporation (Extended Data Fig. 2g,h). By contrast, relative to their respective controls, PAXIP1 depletion in Saos2-dPML cells significantly reduced the number of non-S-phase EdU-positive telomeres. Similar results were also observed in IMR90-ALT#1 cells, where depletion of PAXIP1 alone did not affect cell growth or ALT-associated telomere synthesis, but loss of PAXIP1 in PML-knockout IMR90-ALT#1 cells (dPML#1 or dPML#2) significantly reduced clonogenic growth capacities and non-S-phase telomeric EdU incorporation (Extended Data Fig. 2i-l). Together, these results indicate that PAXIP1-PAGR1 promotes ALT-associated telomere maintenance independently of PML.

### PAXIP1 truncating mutant compensates PML loss in U2OS cells

Our identification of a *PAXIP1* truncating mutation in U2OS cells coincides with its *PML* knockout-induced initial telomere attrition. However, TRF analysis revealed that telomeres in U2OS-dPML cells stopped shortening after 250 population doublings (PD) (Fig. 3a). This seemly recovery of telomere length maintaining capacity prompted us to investigate the underlying mechanism. While western blot analysis confirmed persistent loss of PML in U2OS-dPML cells even after 450 PDs, examination of PAXIP1 revealed a markedly increased abundance of the mutant PAXIP1 protein and also an emergence of a wild-type-sized PAXIP1 band (Fig. 3b). Sanger sequencing of PCR-amplified genomic DNA from U2OS-dPML#1 and dPML#2 cells excluded the possibility of any *PAXIP1* reversion mutation (Extended Data Fig. 3a). Transduction of PAXIP1-targeted sgRNA depleted both PAXIP1 species in U2OS-dPML cells (Extended Data Fig. 3b), further ruling out any non-specific antibody artifact.

**Fig. 3.**
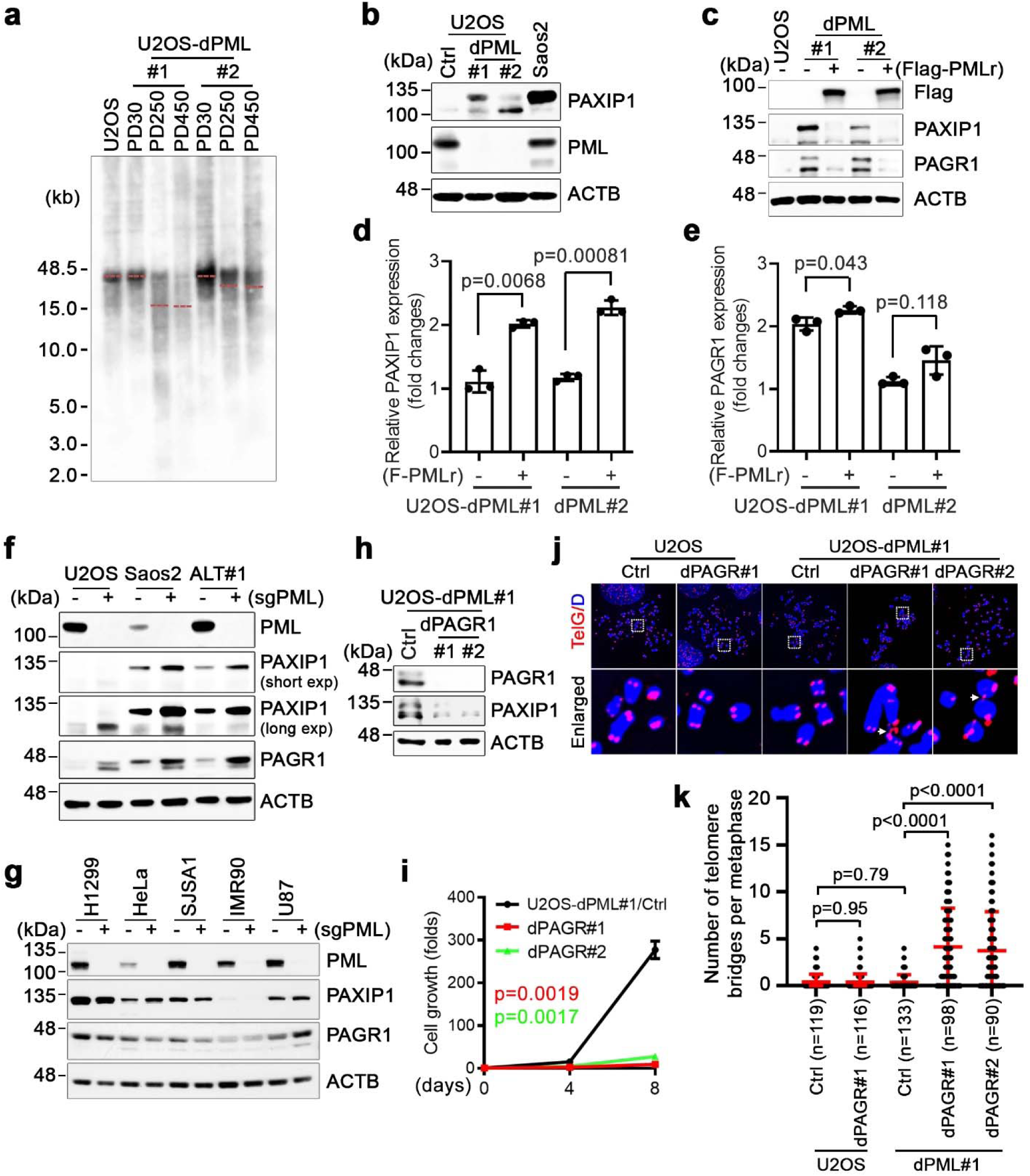
PML regulates PAXIP1-PAGR1 protein abundance. **a,** TRF analysis of telomere length in parental U2OS cells and dPML clones (#1 and #2) at the indicated population doublings (PDs). Genomic DNA was assayed with a digoxigenin (DIG)-labeled TelG probe. **b,** Western blot analysis of PAXIP1 and PML protein expression in parental U2OS cells, U2OS-dPML clones (#1 and #2), and Saos2 cells. **c,** Western blot analysis of Flag, PAXIP1 and PAGR1 protein expression in parental U2OS cells and dPML clones (#1 and #2) transduced with either vector control (−) or CRISPR-resistant, Flag-tagged PML (Flag-PMLr) (+). **d,** RT-qPCR analysis of *PAXIP1* mRNA expression in parental U2OS cells and dPML clones (#1 and #2) transduced with either vector control (−) or Flag-PMLr (+). mRNA levels were normalized to *GAPDH*. **e,** RT-qPCR analysis of *PAGR1* mRNA expression in parental U2OS cells and dPML clones (#1 and #2) transduced with either vector control (−) or Flag-PMLr (+). **f,** Western blot analysis of PML, PAXIP1, and PAGR1 protein expression in U2OS, Saos2, and IMR90-ALT#1 (ALT#1) cells transduced with either sgCtrl (−) or sgPML (+). **g,** Western blot analysis of PML, PAXIP1, and PAGR1 protein expression in the telomerase-positive cell lines H1299, HeLa, SJSA1, IMR90, and U87 transduced with either sgCtrl (−) or sgPML (+). **h-j,** Western blot analysis of PAXIP1 and PAGR1 protein expression (**h**), cell proliferation assays (**i**), and representative telomere FISH images of metaphase spreads (**j**) from parental U2OS-dPML#1 control cells and CRISPR/Cas9-derived PAGR1 deletion clones (dPAGR#1 and #2). Inter-chromosomal telomere fusion events are highlighted by arrows in the enlarged images in (**j**). **k,** Quantification of (**j**), showing the number of telomere fusion events per metaphase. Data represent mean ± s.e.m. from three independent metaphase spread experiments. Statistical analysis was performed using an unpaired two-tailed Student’s t test; P values are shown.

SUMOylation has been implicated in supporting ALT telomere maintenance independently of PML ^31, 43^. To test whether the mutant PAXIP1 is covalently modified by SUMO, we performed immunoprecipitation (IP) of endogenous SUMO2/3 under a denaturing condition in control and U2OS-dPML cells. Immunoblot analysis revealed a ladder of PAXIP1-reactive bands in SUMO2/3 pull-downs from U2OS-dPML#1 and dPML#2, but not from parental U2OS cells (Extended Data Fig. 3c), suggesting that PML loss promotes SUMOylation of mutant PAXIP1 protein and its concomitant stabilization. Consistently, reconstitution of U2OS-dPML#1 and dPML#2 cells with CRISPR-resistant Flag-PMLr markedly reduced the abundance of mutant PAXIP1 and its partner PAGR1 proteins, to levels comparable to those seen in parental U2OS cells (Fig. 3c). Notably, quantitative PCR with reverse transcription (RT-qPCR) analysis indicated that Flag-PMLr re-expression in U2OS-dPML cells did not diminish, but rather elevated, PAXIP1 and PAGR1 mRNA expression relative to their respective controls (Fig. 3d,e), suggesting that PML reconstitution reduces PAXIP1 and PAGR1 abundance primarily via post-transcriptional mechanisms.

We next investigated whether PML regulates PAXIP1 and PAGR1 abundance in ALT cells. Similar to the U2OS-dPML clones that exhibited enriched mutant PAXIP1 protein, acute CRISPR/Cas9-mediated PML depletion in parental U2OS cells resulted in a pronounced increase in both mutant PAXIP1 and PAGR1 protein levels (Fig. 3f). Moreover, PML depletion in Saos2 and IMR90-ALT#1 cells, which express wild-type PAXIP1, similarly elevated the abundance of both PAXIP1 and PAGR1 proteins (Fig. 3f). RT-qPCR again indicated that the increased protein expression was not due to transcriptional regulation (Extended Data Fig. 3d,e), suggesting a form of compensatory posttranslational regulation between PAXIP1-PAGR1 complex and PML. Importantly, this interplay is specific to ALT-positive cells, as PML depletion in non-ALT cell lines (H1299, HeLa, SJSA1, IMR90, or U87) failed to augment PAXIP1 or PAGR1 protein levels (Fig. 3g).

We next asked whether the truncated PAXIP1 mutant in U2OS cells contributes functionally to telomere maintenance. As expected, CRISPR/Cas9-mediated disruption of the mutant *PAXIP1* in parental U2OS cells did not affect the cell growth (Extended Data Fig. 3f,g). By contrast, depletion of the same mutant PAXIP1 in U2OS-dPML cells strongly impaired cell proliferation (Extended Data Fig. 3h,i), suggesting that the truncated PAXIP1 retains substantial biological activity and functions as a hypomorphic allele. In agreement, while PAGR1 - the functional partner of PAXIP1 - was dispensable for parental U2OS controls (Extended Data Fig. 3j,k), its depletion in U2OS-dPML cells greatly impaired the cell proliferation (Fig. 3h,i). Telomere FISH analysis of metaphase spreads from PAGR1-deleted U2OS-dPML cells further revealed a robust increase in inter-chromosomal telomere bridges, chromosome breaks, and chromosomal fragmentation (Fig. 3j,k), indicating defective telomere repair. Together, these findings further support that the PAXIP1-PAGR1 complex and PML drive two parallel pathways that complement one another to promote ALT-mediated telomere maintenance. Each pathway alone is sufficient to sustain ALT cell survival, but simultaneous disruption of both results in severe ALT telomere repair defects and a synthetic lethal phenotype.

### STAG2 acts epistatically with PAXIP1-PAGR1 in ALT telomere maintenance

The PAXIP1-PAGR1 heterodimer is reportedly a constituent of the KMT2D/C methyltransferase complex that has been implicated in transcriptional regulation ^38, 44, 45^. However, RNA sequencing analysis revealed that depletion of PAXIP1 in parental Saos2 cells produced no significant changes in gene expression (Extended Data Fig. 4a). In Saos2-dPML cells, PAXIP1 depletion resulted in 224 differentially expressed genes (134 genes downregulated and 90 upregulated relative to controls); however, no changes were observed in genes associated with DNA repair or telomere maintenance (Extended Data Fig. 4b). Gene set enrichment analysis (GSEA) identified significant enrichments of gene sets such as MYC-targets, E2F-targets, and NF-κB signaling (Extended Data Fig. 4c), changes that likely reflect secondary effects linked to cell stress and/or impaired cell proliferation.

To pinpoint the functional network surrounding PAXIP1 and PAGR1, we next analyzed Pearson correlation coefficients of gene dependency scores from the DepMap database (https://depmap.org) ^46, 47^. This analysis revealed a strong co-dependency between PAXIP1 and PAGR1 themselves and also a pronounced correlation between both proteins with multiple cohesin components, particularly the cohesin subunit STAG2 (Extended Data Fig. 5a,b). Cohesin complexes in vertebrate somatic cells carry one of two STAG isoforms, STAG1 or STAG2 ^48^. While the two paralogs share overlapping functions throughout the cell cycle, they also perform distinct roles in genome organization and gene regulation ^49^. Particularly, STAG1-containing cohesin, but not STAG2-cohesin, is uniquely required for telomeric cohesion formation ^50–52^, whereas STAG2-cohesin has been implicated in DNA damage repair and replication fork progression ^53–55^. To determine whether STAG2-cohesin participates in ALT-mediated telomere maintenance, we examined its telomeric association in U2OS, Saos2, and IMR90-ALT#1 cells. Chromatin immunoprecipitation (ChIP) followed by telomere dot-blot analysis revealed robust STAG2 enrichment at telomeres in both Saos2 and IMR90-ALT#1 cells, whereas telomeric signals from U2OS ChIP samples were below detectable levels (Fig. 4a). This pattern closely correlates with PAXIP1 and PAGR1 protein expression in these cell lines (Fig. 4b).

**Fig. 4.**
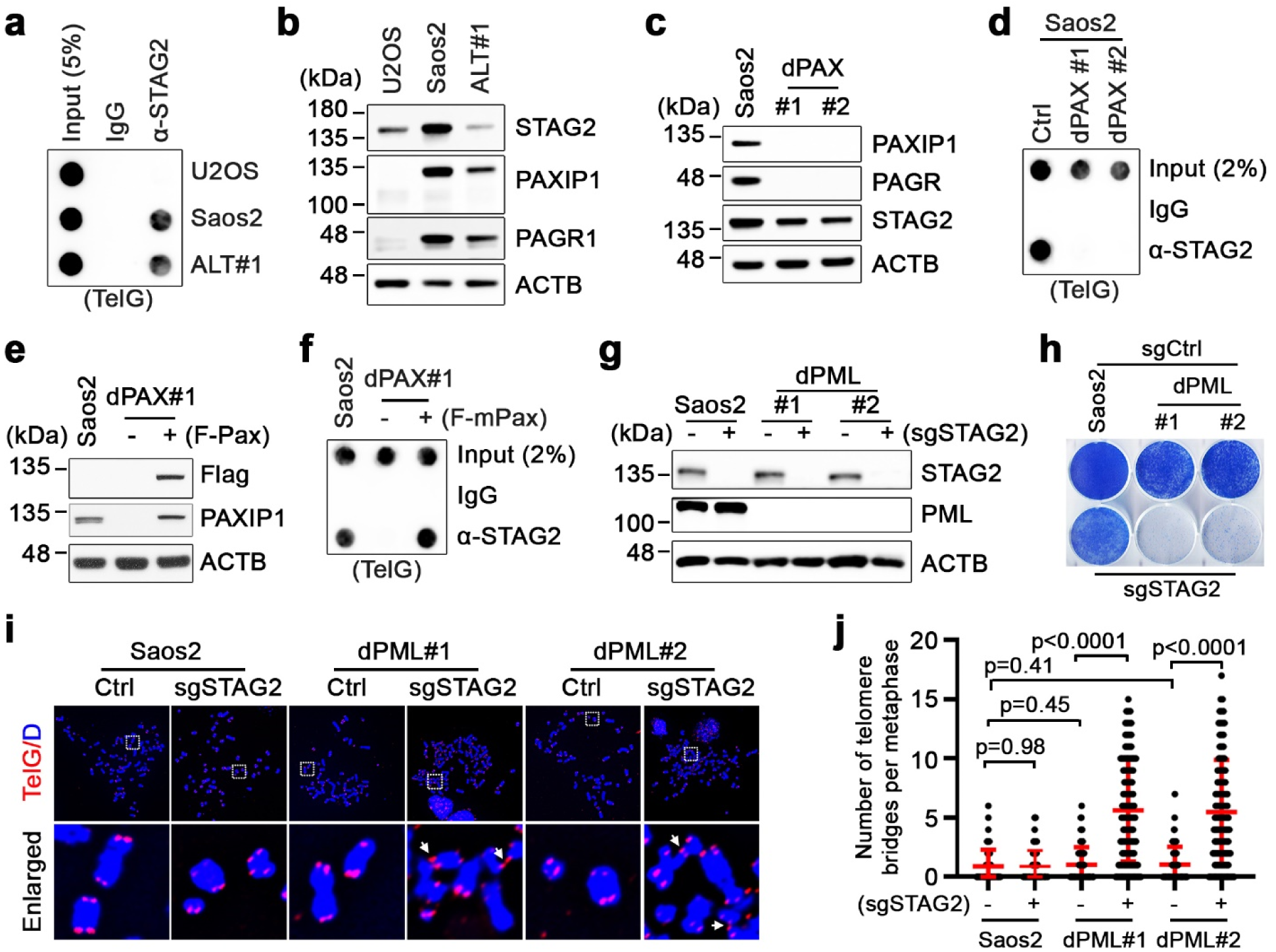
PAXIP1-PAGR1 directs STAG2-cohesin recruitment to promote ALT telomere maintenance. **a,** Telomere dot-blot analysis of anti-STAG2 ChIP in U2OS, Saos2, and IMR90-ALT#1 (ALT#1) cells. IgG ChIP was included as a control for non-specific signals. Input and ChIP DNA were hybridized with a DIG-labeled TelG probe. **b,** Western blot analysis of STAG2, PAXIP1, and PAGR1 protein expression in U2OS, Saos2, and IMR90-ALT#1 (ALT#1) cells. ACTB served as a loading control. **c,** Western blot analysis of PAXIP1, PAGR1, and STAG2 protein expression in parental Saos2 cells and two CRISPR/Cas9-derived PAXIP1 deletion clones (dPAX#1 and #2). **d,** Telomere dot-blot analysis of anti-STAG2 ChIP in parental Saos2 cells and Saos2-dPAX clones (#1 and #2). **e,** Western blot analysis of ectopically expressed Flag-mPax (detected with anti-Flag antibody) and total PAXIP1 protein expression in parental Saos2 cells and Saos2-dPAX#1 clone transduced with vector (−) or Flag-mPax (+). **f,** Telomere dot-blot analysis of anti-STAG2 ChIP in parental Saos2 cells and Saos2-dPAX#1 clone transduced with vector (−) or Flag-mPax. **g,** Western blot analysis of STAG2 and PML protein expression in parental Saos2 cells and Saos2-dPML clones (#1 and #2) following transduction with sgCtrl (−) or sgSTAG2 (+). **h,** Crystal violet-based clonogenic growth assay of parental Saos2 cells and Saos2-dPML clones transduced with sgCtrl (−) or sgSTAG2 (+). Crystal violet staining was conducted on day 17 post-seeding. **i,** Representative telomere FISH images of metaphase spreads from parental Saos2 cells and dPML clones (#1 and #2) transduced with sgCtrl or sgSTAG2. Inter-chromosomal telomere fusion events are highlighted by arrows in the enlarged images. **j,** Quantification of (**i**), showing the number of telomere fusion events per metaphase in parental Saos2 cells and dPML clones transduced with sgCtrl or sgSTAG2. Data represent mean ± s.e.m. from three independent metaphase spread experiments. Statistical analysis was performed using an unpaired two-tailed Student’s t test; P values are shown.

PAXIP1 has previously been implicated in regulating chromatin cohesin stability ^56^. To test whether PAXIP1 is required for STAG2-cohesin recruitment to ALT telomeres, we generated clonal Saos2 derivatives carrying CRISPR/Cas9-mediated *PAXIP1* deletions (Fig. 4c). Analysis of anti-STAG2 ChIP samples by telomere dot blot demonstrated that loss of PAXIP1 completely abolished STAG2-cohesin recruitment to telomeres in Saos2-dPAX cells (Fig. 4d). Reintroduction of an sgRNA-resistant Flag-mPax transgene into Saos2-dPML cells restored telomeric STAG2 recruitment (Fig. 4e,f). Similar defects in telomeric STAG2-cohesin loading also occurred in *PAGR1*-deleted Saos2-dPAGR cells (Extended Data Fig. 5c,d), indicating that the PAXIP1-PAGR1 complex is instrumental in driving STAG2-cohesin recruitment to ALT telomeres.

Our findings indicate that PAXIP1-PAGR1 and PML complement one another to promote ALT-mediated telomere maintenance. To assess whether PAXIP1-PAGR1 acts through STAG2 to fulfill this function, we transduced Saos2 and Saos2-dPML cells with sgRNA targeting *STAG2* (Fig. 4g). Similar to PAXIP1 and PAGR1, STAG2 was largely dispensable for parental Saos2 cell growth. In contrast, depletion of STAG2 in Saos2-dPML cells phenocopied the effects of PAXIP1 or PAGR1 loss, resulting in a pronounced reduction in clonogenic growth (Fig. 4h) and a substantial increase in metaphase formation of inter-chromosomal telomere bridges (Fig. 4i,j). Consistently, sgRNA-mediated STAG2 depletion also significantly impaired the clonogenic growth of IMR90-ALT#1-dPML cells (Extended Data Fig. 5e,f), suggesting that STAG2 acts epistatically with the PAXIP1-PAGR1 complex in maintaining ALT telomeres. By comparison, depletion of the other STAG paralog, STAG1 - individually or in combination with PML - failed to suppress clonogenic growth of Saos2 cells (Extended Data Fig. 5g,h), despite its documented involvement in telomeric sister chromatid cohesion ^50–52^.

### PAXIP1-PAGR1 physically interacts with the STAG2-RAD21 cohesin subcomplex

The strong co-dependency and functional epistasis observed between PAXIP1-PAGR1 and STAG2 prompted us to investigate whether these proteins physically interact. To this end, we co-expressed in insect cells the following: human PAGR1, the PAGR1-binding region of PAXIP1 (amino acids 94-183, encompassing BRCT domain I/II), human STAG2, and the STAG2-binding region of RAD21 (amino acids 281-420). Co-purification confirmed the formation of a four-protein complex (Extended Data Fig. 6a,b). Extensive sample screening followed by data collection on a 300-kV Krios G4 cryo-EM yielded a “Z”-shaped structure at 2.49 Å resolution (Extended Data Fig. 6c-g and Extended Data Table 1). The final 3D model included STAG2 residues 87-250, 262-440, 452-837, 851-961 and 1002-1008; RAD21 residues 322-398; and PAGR1 residues 164-172 (Fig. 5a,b). Notably, the cryo-EM density for PAXIP1 (BRCT I/II) was not well resolved, presumably due to structural flexibility.

**Fig. 5.**
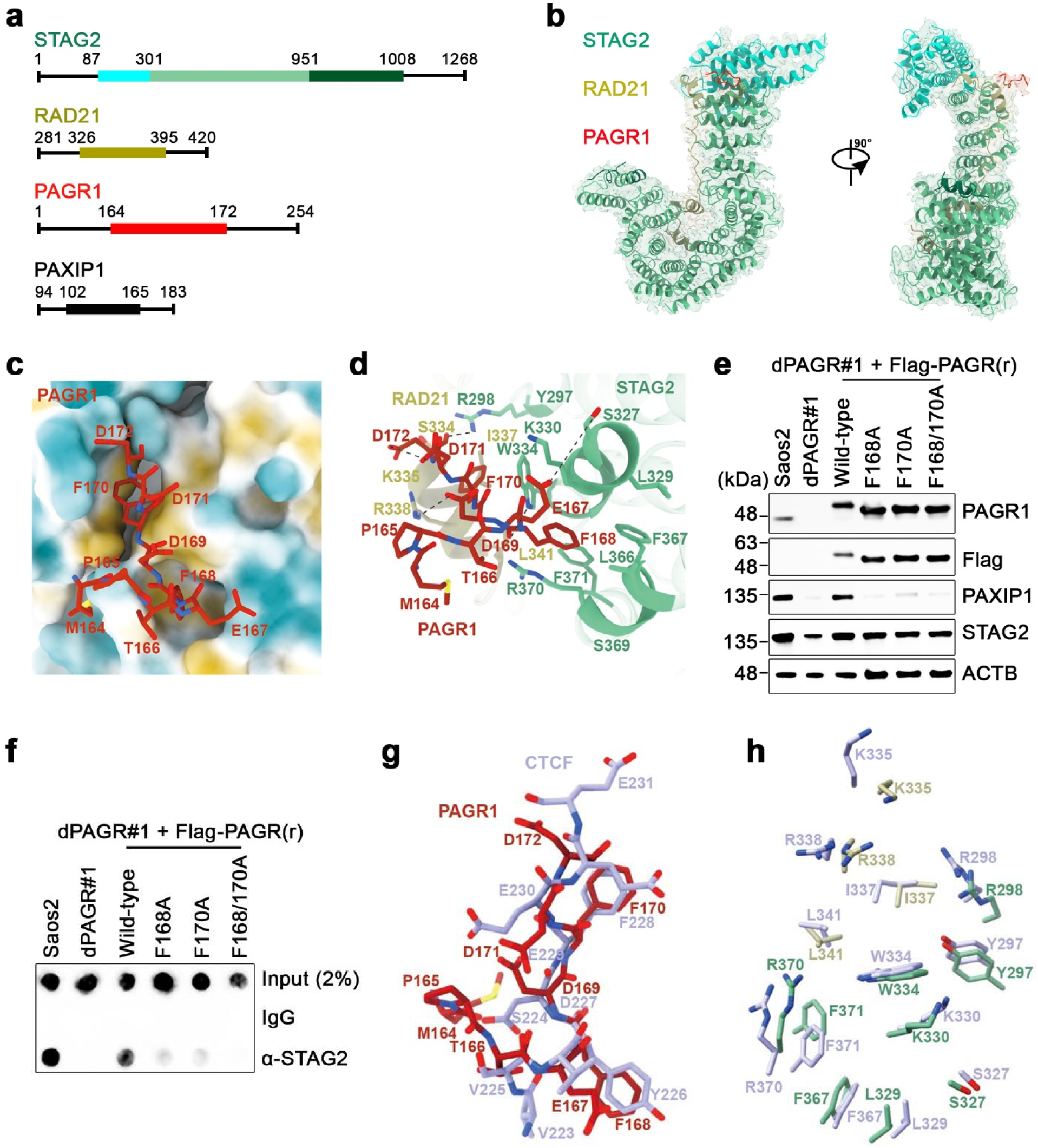
PAXIP1-PAGR1 physically interacts with the STAG2-RAD21 cohesin subcomplex. **a,** Schematic representation of the STAG2, RAD21, PAGR1, and PAXIP1 expression constructs used in this study. The regions shown in the electron density map are highlighted by respective colored boxes. **b,** Cartoon illustrations of the PAGR1-STAG2-RAD21 structure. STAG2, RAD21, and PAGR1 are colored in green, orange, and red, respectively. **c,** Hydrophobic pockets of STAG2-RAD21 cohesin subcomplex engage with PAGR1. Residues F168 and F170 of PAGR1 bind to the respective hydrophobic pockets formed jointly by STAG2 and RAD21. **d,** Interaction interfaces of PAGR1 with STAG2 and RAD21. Black dashed lines indicate hydrogen bonds formed at the interfaces. **e,** Western blot analysis of total PAGR1, ectopically expressed CRISPR-resistant Flag-tagged PAGR1 (Flag-PAGRr, detected with anti-Flag antibody), PAXIP1, and STAG2 protein expression in parental Saos2 cells, a PAGR1 deletion clone (dPAGR#1), and dPAGR#1 cells transduced with indicated wild-type or mutant Flag-PAGRr constructs. ACTB is used as a loading control. **f,** Telomere dot-blot analysis of anti-STAG2 ChIP in parental Saos2, dPAGR#1 cells, or dPAGR#1 clones transduced with the indicated wild-type or mutant Flag-PAGRr constructs. Input and ChIP DNAs were hybridized with a DIG-labeled TelG probe. **g,** Superimposed structures of proteins PAGR1 and CTCF. **h,** Conformational changes at the STAG2-RAD21 interface. Superposition of PAGR1-bound and CTCF-bound (PDB 6QNX) STAG2-RAD21 structures reveals distinct conformational rearrangements of key interface residues.

STAG2 comprises an N-terminal domain, a central HEAT repeats domain, and a C-terminal domain. While RAD21 interacts along one side of STAG2, spanning from the N-terminal to the C-terminal domain, PAGR1 engages through its conserved FDF motif to a hydrophobic interface that is formed jointly by STAG2 and RAD21 (Fig. 5c). Specifically, PAGR1 residues F168 and F170 occupy two distinct hydrophobic cavities: F168 inserts into a pocket formed by STAG2 L366, F367, and F371 (cavity 1); F170 occupies the second cavity lined by STAG2 Y297, R298, W334, and RAD21 K335, I337, and R338 (cavity 2) (Fig. 5d). Additional interactions contribute to binding specificity and stability - hydrogen bonds are formed between PAGR1 residues D172, F168, E167 and STAG2 residues R298, W334, and S327, respectively. Furthermore, PAGR1 residues D172 and D169 establish salt bridges with RAD21 residues K335 and R338, providing additional anchoring contact.

To quantify the binding affinity, we purified PAXIP1 (94-183), PAGR1, STAG2, and RAD21 (281-420) separately. Biolayer interferometry (BLI) assays revealed that PAXIP1-PAGR1 binds the STAG2-RAD21 cohesin subcomplex at a KD of 232.7 ± 12 nM (Extended Data Fig. 7a). To validate functional importance of the FDF motif, we reconstituted Saos2-dPAGR cells with sgRNA-resistant, Flag-tagged wild-type PAGR1 or mutants carrying alanine substitution at residues F168 and/or F170 (Fig. 5e). Telomere dot-blot analysis of anti-STAG2 ChIP samples demonstrated that wild-type PAGR1 - but not the F168A, F170A, or F168/170A mutants - restored telomeric STAG2-cohesin recruitment in Saos2-dPAGR cells (Fig. 5f), confirming F168 and F170 of the FDF motif as critical determinants of PAGR1 binding to STAG2-cohesin. Consistently, expression of sgRNA-resistant wild-type PAGR1, but not STAG2-binding-defective mutants, rescued the growth defect caused by PAGR1 depletion in Saos2-dPML cells (Extended Data Fig. 7b,c).

Notably, the composite PAGR1-binding interface formed by STAG2 and RAD21 overlaps with the reported interaction surface utilized by CTCF ^57^ and SGO1 ^58^, suggesting a potentially conserved mode of cohesion regulation. To compare these interactions, we superimposed the reported CTCF-bound structure (Extended Data Fig. 7d) ^57^ onto our PAGR1-STAG2-RAD21 complex. This analysis revealed remarkable correspondence. Specifically, PAGR1 residues T166, E167, F168, D169, and F170 align closely with CTCF residues S224, V225, Y226, D227 and F228, respectively (Fig. 5g). However, PAGR1 residues M164, P165, D171, and D172 exhibit conformations distinct from those of the corresponding CTCF residues V223, E229, E230 and E231. Moreover, several STAG2 residues, including K335, R338, R370, D326 and R298, also adopt different conformations in the two complexes (Fig. 5h), likely reflecting their distinct regulatory mechanisms and biological functions.

### PAXIP1-PAGR1 directs cohesin recruitment during break-induced telomere repair

Previous studies have shown that STAG1-containing cohesin, but not STAG2-cohesin, is specifically required for cohesion establishment at telomeres ^50–52^. In agreement, telomere dot-blot analysis of telomerase-immortalized IMR90-T cells revealed undetectable telomeric STAG2-cohesin association (Fig. 6a), correlating with the absence of telomeric DNA repair (Extended Data Fig. 8a,b) and low abundance of PAXIP1 and PAGR1 proteins (Fig. 6b). By contrast, analysis of isogenic ALT-positive IMR90-ALT cells showed robust telomeric STAG2 enrichment with markedly higher telomere DNA repair and PAXIP1 and PAGR1 protein expression relative to IMR90-T cells, consistent with our findings that the PAXIP1-PAGR1 complex is crucial for STAG2-cohesin recruitment to ALT telomeres.

**Fig. 6.**
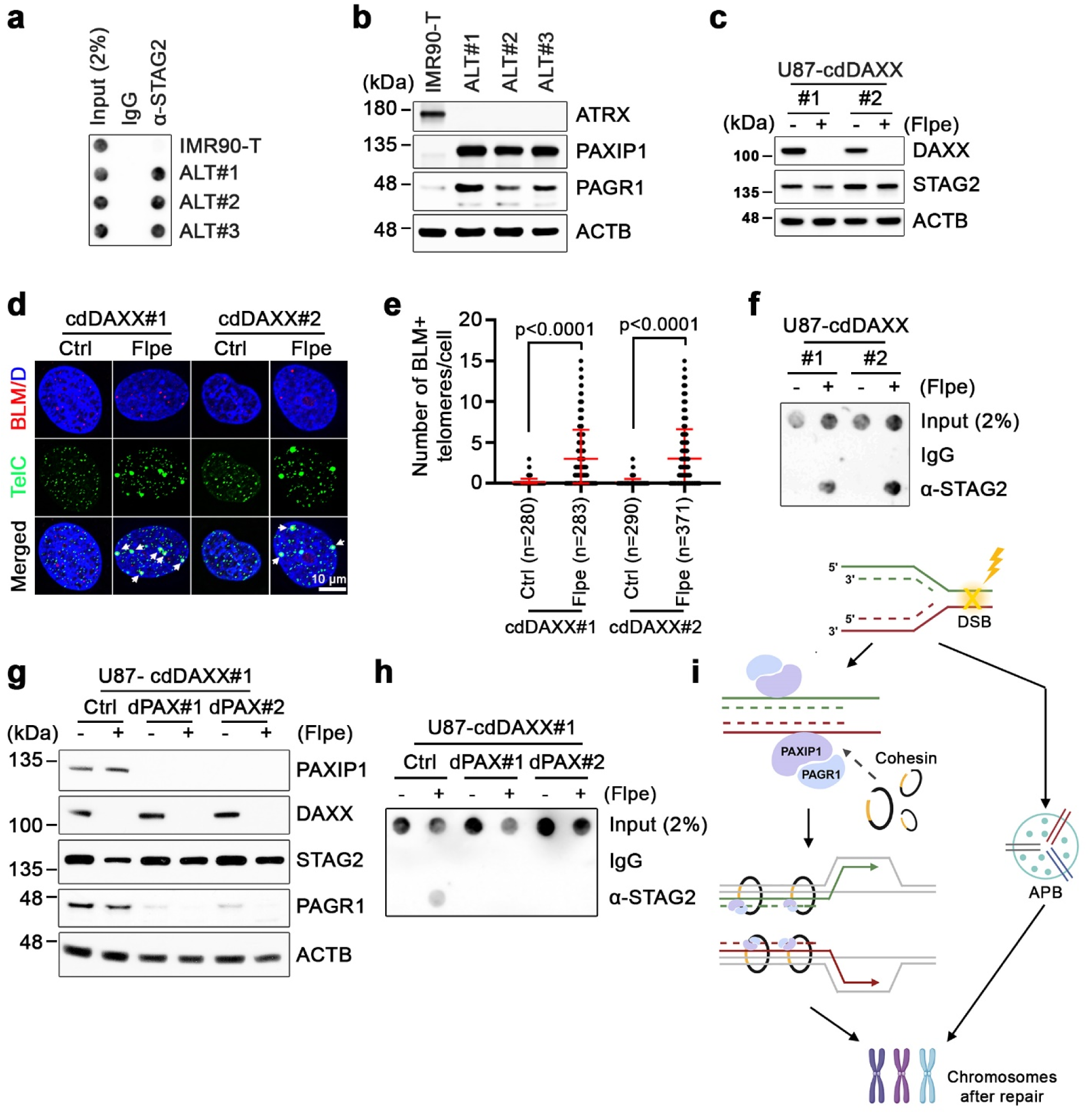
The PAXIP1-PAGR1 complex controls telomere break-induced de novo STAG2-cohesin recruitment. **a,** Telomere dot-blot analysis of anti-STAG2 ChIP in IMR90-T, IMR90-ALT#1, ALT#2, and ALT#3 cells. IgG ChIP was included as a control for non-specific signals. Input and ChIP DNAs were hybridized with a DIG-labeled TelG probe. **b,** Western blot analysis of ATRX, PAXIP1, and PAGR1 protein expression in IMR90-T, IMR90-ALT#1, ALT#2, and ALT#3 cells. ACTB serves as a loading control. **c,** Western blot analysis of DAXX and STAG2 protein expression in telomerase-positive U87-cdDAXX cells (#1 or #2) transduced with either control (−) or Flpe construct (+). **d,** Representative IF-FISH images showing BLM colocalization with telomeres (TelG) in G2/M phase-synchronized U87-cdDAXX cells (#1 or #2) transduced with either control (−) or Flpe construct (+). Arrows denote BLM colocalized telomere foci. **e,** Quantification of (**d**), showing the number of BLM-positive telomere foci per cell in G2/M-synchronized U87-cdDAXX cells (#1 or #2) transduced with either control (−) or Flpe construct (+). Data represent mean ± s.e.m. from three independent experiments. Statistical analysis was performed using an unpaired two-tailed Student’s t test; P values are shown. **f,** Telomere dot-blot analysis of anti-STAG2 ChIP in U87-cdDAXX cells (#1 or #2) transduced with either control (−) or Flpe construct (+). **g,** Western blot analysis of PAXIP1, DAXX, STAG2, and PAGR1 protein expression in parental U87-cdDAXX#1 cells (Ctrl) or PAXIP1 deletion clones (dPAX#1 and #2) transduced with either control (−) or Flpe construct (+). **h,** Telomere dot-blot analysis of anti-STAG2 ChIP in parental U87-cdDAXX#1 control and dPAX clones (#1 and #2) transduced with either control (−) or Flpe construct (+). **i,** Model depicting how the PAXIP1-PAGR1-cohesin and PML nuclear body-associated pathways complement one another to promote break-induced telomere synthesis and repair.

We next asked whether this observed telomeric STAG2-cohesin recruitment is unique to ALT-positive cells or instead represents a more general response to telomeric DNA damage. To address this question, we utilized a previously established conditional *DAXX* knockout system (cdDAXX), in which endogenous *DAXX* was deleted from telomerase-positive U87 glioblastoma cells that carry a Flpe recombinase-targeted DAXX rescue cassette ^36^. In line with our previous report ^36^, Flpe-mediated *DAXX* deletion in U87-cdDAXX#1 and #2 cells triggered telomere replication stress and subsequent break-induced telomere repair, as evidenced by a marked increase in telomere-associated BLM foci (Fig. 6c-e). ChIP coupled with telomere dot-blot analysis further revealed a robust induction of telomeric STAG2-cohesion establishment following Flpe-mediated DAXX depletion (Fig. 6f). Importantly, control cdDAXX cells exhibited undetectable levels of telomeric STAG2-cohesin, consistent with previous reports that STAG2 is not involved in cohesion establishment of normal telomeres ^51, 52^. These data support the notion that damage-induced cohesion is established mainly through *de novo* recruitment rather than repositioning of pre-existing cohesin complexes.

We next examined whether the telomeric DNA damage-induced STAG2-cohesin recruitment relies on the PAXIP1-PAGR1 complex. To this end, we generated clonal *PAXIP1* deletion (dPAX) derivatives of U87-cdDAXX cells (Fig. 6g). Whereas Flpe-mediated DAXX depletion in parental cdDAXX#1 cells induced robust telomeric STAG2 recruitment, knockout of *PAXIP1* abrogated the damage-induced telomeric STAG2-cohesin enrichment (Fig. 6h). Similar results were observed in *PAGR1*-deleted U87-cdDAXX cells, where loss of PAGR1 completely abolished the damage-induced telomeric STAG2-containing cohesion establishment (Extended Data Fig. 8c,d), indicating that the PAXIP1-PAGR1 complex controls *de novo* establishment of STAG2-containing cohesion during break-induced telomeric DNA repair.

## Discussion

Our findings reveal a dynamic DNA damage response that bridges DSB recognition to the assembly of cohesin capable of executing BIR and damage repair. We propose that PAXIP1 functions as a DSB sensor through its BRCT domains, by recruiting PAGR1 and subsequently STAG2-containing cohesin to sites of DNA damage to promote homology-based BIR, whereas ALT-associated PML nuclear bodies (APBs) independently provide an additional recombinogenic microenvironment to facilitate ALT-mediated repair (Fig. 6i). Our data indicate that these two pathways operate in parallel and complement one another in promoting ALT-directed telomere maintenance. Each pathway alone is sufficient to sustain ALT cell survival, but simultaneous disruption of both leads to profound defects in telomere maintenance and synthetic lethality of ALT-positive cells, underscoring their functional redundancy and collective importance.

Although cohesin recruitment to DSBs to facilitate homology-directed DNA damage repair has been extensively documented in both mammals and yeast ^5–9, 11–13, 15^, how cohesion is established in the context of DNA repair remains less clear. The molecular and structural evidence provided in this study identifies the PAXIP1-PAGR1 complex as a bridge linking the DNA damage response to repair-associated *de novo* cohesion establishment. Our data further reveal that physical interaction of STAG2-RAD21 cohesin subcomplex with the PAXIP1-PAGR1 complex is essential for damage-induced cohesion establishment at telomeres. These observations suggest two possible mechanistic models. In the simplest scenario, the PAXIP1-PAGR1 complex acts as a cohesin-loading factor that directly recruits cohesin to sites of damage. Alternatively, PAXIP1-PAGR1 may primarily regulate cohesin retention on damaged chromatin. The identified PAGR1 cohesin binding interface overlaps with a conserved surface utilized by CTCF ^57^ and SGO1 ^58^. Notably, SGO1 binding has been shown to protect cohesin from WAPL-mediated cohesin unloading ^58^. Along this line, it stands to reason that the PAXIP1-PAGR1 complex could similarly function as a cohesion-stabilizing factor by antagonizing WAPL-mediated cohesin release, a key regulatory process for cohesin stability ^59, 60^. These two models are not mutually exclusive. Future measurement of cohesin stability in the context of DNA damage will be necessary to quantify their relative contributions.

Our study demonstrates that the PAXIP1-PAGR1 complex directs *de novo* STAG2-containing cohesion establishment in supporting break-induced telomere damage repair. We envisage that analogous process may operate at other vulnerable genomic loci during BIR and perhaps more broadly, during HR-based repair of conventional two-ended DSBs. Beyond its role in DNA damage repair, cohesin-regulated recombination contributes to a wide range of biological processes that depends on controlled DNA rearrangements ^2^. One notable example is antibody class switch recombination (CSR) ^61, 62^. Interestingly, loss of either STAG2 or PAXIP1-PAGR1 impairs CSR without broadly disrupting DNA damage repair pathways ^38, 61, 63^, raising another scenario possibly involving PAXIP1-PAGR1-cohesin interplay. Taken together, our findings uncover a critical role for the PAXIP1-PAGR1 complex in directing cohesin-regulated DNA recombination, a mechanism likely conserved across diverse biological activities.

## Methods

### Cell lines

The human cell lines HEK293T, HeLa, IMR90, NCI-H1299, Saos2, SJSA1, U2OS, and U87 were obtained from ATCC. The High Five insect cell line was obtained from Thermo Fisher Scientific. IMR90-T, IMR90-ALT#1, #2, #3, and U87-cdDAXX cells were generated from IMR90 or U87 cells as previously described^36^. Clonally derived *PML*-deletion cell lines (HeLa, IMR90-ALT#1, Saos2, U2OS, and U87), *PAXIP1*-deletion cell lines (Saos2, U2OS, and U87-cdDAXX), and *PAGR1*-deletion cell lines (Saos2, U2OS, and U87-cdDAXX) were generated from respective parental cells transduced with lentiCRISPR-v2 encoding sgRNA targeting *PML*, *PAXIP1*, or *PAGR1*, followed by blasticidin selection as previously described^36^. HEK293T, HeLa, IMR90, NCI-H1299, SJSA1, U2OS, U87, and their derived cell lines were cultured in DMEM (Corning) supplemented with 10% FBS (Avantor Seradigm) and 1% penicillin–streptomycin (Corning). Saos2 and its derived cell lines were cultured in DMEM (Corning) supplemented with 15% FBS (Avantor Seradigm) and 1% penicillin–streptomycin (Corning). All cell lines were maintained in a humidified incubator at 37°C with 5% CO2 according to standard protocols. High Five insect cells were grown in Express Five SFM (Thermo Fisher Scientific) supplemented with 18 mM L-glutamine (Thermo Fisher Scientific) in a humidified incubator at 27°C with 5% CO2, following manufacturer’s instructions. All cell lines were confirmed negative for mycoplasma contamination using the MycoAlert Plus Mycoplasma Detection Kit (Lonza).

### sgRNA cloning and gene deletion

All sgRNAs were cloned into lentiCRISPRv2 blast (Addgene plasmid #98293), LRPuro^36^ (U6-sgRNA-Puro, reconstructed from LRG2.1 - a gift from C. Vakoc; Addgene plasmid #108098), LRNeo^36^ (U6-sgRNA-Neo, reconstructed from LRPuro), and LRBlast^36^ (U6-sgRNA-Blast, reconstructed from LRPuro). Single sgRNAs were generated by annealing two DNA oligos and ligating them into the indicated BsmB1-digested vector.

For single-gene deletion, indicated cells were infected with lentiCRISPRv2 blast targeting the gene of interest and selected with blasticidin. For double-gene deletion, cells already expressing lentiCRISPRv2 blast targeting the first gene were further infected with LRPuro, LRNeo, or combination thereof targeting the second gene, followed by selection with the appropriate antibiotics. A complete list of sgRNA sequences is provided in Supplementary Table 1.

### cDNA expression constructs and lentivirus production

For cDNA expression experiments, full length cDNAs were cloned into the lentiviral expression vectors pLU-IRES-Puro, -Blast, -Neo, or Hygro, each containing an N-terminal 3xFlag tag. CRISPR sgRNA-resistant synonymous mutations and functional domain point mutations were introduced by PCR mutagenesis using the NEBuilder HiFi DNA Assembly Master Mix (E2631, NEB). Stable cell lines were generated using the pLU vectors and selected with appropriate antibiotics.

Lentiviruses were produced by co-transfecting the indicated plasmids along with packaging vectors into HEK293T packaging cells, as previously described ^36^. Briefly, to generate lentivirus, 8 × 10^6^ 293T cells in 100 mm tissue culture dishes were transfected with a mixture containing 8.5 μg of plasmid DNA, 4 μg of pMD2.G, 6 μg of psPAX2 packaging vectors, and 45 μl of 1 mg/mL Polyethylenimine (PEI 25000). The media was replaced 6-8 h post transfection. Virus-containing supernatant was collected at 48 and 72 hours post-transfection and pooled.

For infection, virus-containing supernatant was mixed with the indicated cell lines supplied with 8 μg/mL polybrene and centrifuged at 2,000 rpm for 30 min at room temperature. Fresh media was added 24 hours post-infection. When selection was required, antibiotics (10 μg/mL blasticidin, 2 μg/mL puromycin, 500 μg/mL G418, and/or 200 μg/mL hygromycin) were added 48 hours post-infection.

### Western blot analysis

Cells were lysed in RIPA buffer (150 mM NaCl, 50 mM Tris, 0.5% Na-Deoxycholate, 0.1% SDS and 1% NP-40). Equal amounts of protein were resolved by electrophoresis on Nupage Novex 4-12% Bis-Tris Gel (Thermo Fisher Scientific) as previously described ^36^. Primary antibodies used were as follows: β-ACTIN (ACTB) (1:5,000; #2228, Sigma), ATRX (1:1,000; HPA001906, Sigma-Aldrich), DAXX (1:1,000; D7810, Sigma-Aldrich), Flag (1:1,000; F1804, Sigma), PML (1:500; sc-966, Santa Cruz Biotechnology), PAXIP1 (1:1,000; ab70434, abcam), PAGR1 (1:1,000; ABE1863, Millipore-Sigma), STAG1 (1:1,000; A300-156A, Fortis Life Sciences), STAG2 (1:1,000; #4239, Cell Signaling), STAG2 (1:2,000; ab4464, abcam), TUBULIN (1:2,000; ab15246, abcam). Secondary antibodies used were donkey anti-rabbit HRP (1:1,000; sc-2077, Santa Cruz Biotechnology), donkey anti-mouse HRP (1:1,000; sc-2096, Santa Cruz Biotechnology), and donkey anti-goat HRP (1:1,000; sc-2056, Santa Cruz Biotechnology). Antibody signals were detected using the SuperSignal™ West Pico or Femto Chemiluminescent substrate (Thermo Fisher Scientific). High-resolution digital images were captured using the Azure Biosystems Imaging System.

### Immunofluorescence and fluorescence in situ hybridization (IF-FISH)

Indirect immunofluorescence (IF) combined with fluorescence *in situ* hybridization (FISH) analysis was performed as previously described^36^. Briefly, cells grown on circular coverslips were cooled on ice, rinsed once with 1x PBS, and placed in pre-extraction buffer (0.1% Triton X-100, 20 mM HEPES-KOH pH 7.9, 50 mM NaCl, 3 mM MgCl2, 300 mM sucrose) for 5 min on ice. The coverslips were then rinsed once with 1x PBS, fixed with 4% paraformaldehyde for 15 min at room temperature, followed by cold methanol (−20°C) for 10 min, and permeabilized in PBS containing 0.5% Triton X-100 for 10 min at room temperature. After washing with 1x PBS, coverslips were incubated for 60 min in blocking solution (1% BSA, 10% FBS, 0.2% fish gelatin, 0.1% Triton X-100, 1 mM EDTA in 1x PBS) prior to immunostaining. Primary antibodies were diluted in blocking solution as follows: BLM (1:500; A300-110A, Fortis Life Sciences), Flag (1:200; F1804, Sigma-Aldrich), and PML (1:200; sc-966, Santa Cruz Biotechnology). Following incubation with primary antibodies for 2 hours at room temperature, cells were washed three times with PBST (1xPBS containing 0.1% Tween-20) and then incubated with fluorophore-conjugated secondary antibodies diluted in blocking solution for 45 min at room temperature. After three washes with PBST, cells were fixed again with 4% paraformaldehyde for 10 min, washed in 1x PBS, dehydrated through ethanol series (70%, 95%, 100%), and air-dried. Coverslips were denatured for 10 min at 85°C in hybridization mix (70% formamide, 10 mM Tris-HCl, pH 7.2, and 0.5% blocking solution [Roche]) containing 100nM telomeric PNA probe TelC-FITC (F1009, PNA Bio), followed by hybridization for 2 hours at room temperature in dark, moisturized chambers. Coverslips were washed three times with Wash Solution (70% formamide, 2x SSC) for 10 min each and in 2x SSC, 0.1% tween-20 three times for 5 min each, before mounting with Vectashield mounting medium containing DAPI (Vector Labs). Images were captured using a 60x lens on an Olympus FLUOVIEW laser scanning confocal microscope or a 60x lens on a Nikon Eclipse Ti2-E live-cell and Deconvolution Imaging System.

### Cell cycle synchronization

Cells were synchronized in G1/S phase by thymidine treatment, and in G2/M phase by sequential treatment with thymidine followed by CDK1 inhibitor Ro-3306 (S7747, Selleckchem). Briefly, for G1/S synchronization, cells were cultured in medium containing 2 mM thymidine (Sigma-Aldrich) for 16-24 hours. For G2/M synchronization, G1/S-synchronized cells were washed twice with PBS and once with growth media, then released into fresh medium for 3 hours, followed by treatment with 10 µM CDK1 inhibitor for 12 hours.

### Detection of non-S-phase telomere DNA synthesis (EdU labeling)

To visualize non-S phase telomere DNA synthesis, G2/M-synchronized cells were incubated with 50 µM EdU for 2 h. Cells were then permeabilized and fixed with 4% paraformaldehyde in PBS. The Click-iT® Alexa Fluor 594 azide reaction was performed according to the manufacturer’s instructions (C10339, Thermo Fisher Scientific). After washing with 1x PBS, cells were incubated for 60 min in blocking solution (1% BSA, 10% FBS, 0.2% fish gelatin, 0.1% Triton X-100, 1 mM EDTA in 1x PBS), followed by immunostaining with an anti-PML antibody for 2 hours at room temperature. After secondary antibody incubation and three washes with PBST, cells were fixed again with 4% paraformaldehyde for 10 min at room temperature, washed in 1x PBS, dehydrated through ethanol series (70%, 95%, 100%), and air-dried. Coverslips were denatured for 10 min at 85°C in hybridization mix (70% formamide, 10 mM Tris-HCl, pH 7.2, and 0.5% blocking solution [Roche]) containing 100 nM telomeric PNA probe TelC-FITC (F1009, PNA Bio), followed by hybridization for 2 hours at room temperature in dark, moisturized chambers. Coverslips were washed three times with Wash solution (70% formamide, 2x SSC) for 10 min each, and three times with 2x SSC containing 0.1% tween-20 for 5 min each. After the final washes, coverslips were mounted using Vectashield mounting medium with DAPI (Vector Labs). Images were captured with a 60x lens on an Olympus FLUOVIEW laser scanning confocal microscope or a 60x lens on a Nikon Eclipse Ti2-E live-cell and Deconvolution Imaging System.

### Terminal restriction fragment (TRF) analysis

TRF analysis was conducted as previously described^36^. Briefly, genomic DNA was extracted using the DNeasy Blood & Tissue kit (69506, Qiagen) according to the manufacturer’s instructions. For telomere length analysis, approximately 5 μg genomic DNA was digested with *Alu*I and *Mbo*I restriction endonucleases, fractionated on a 0.7% agarose gel, denatured, and transferred onto a GeneScreen Plus hybridization membrane (PerkinElmer). The membrane was cross-linked and hybridized with either 5′-end-labeled ^32^P-TelG probe or a digoxigenin (DIG)-labeled DIG-TelG probe.

For detection with ^32^P-TelG probe, the membrane was incubated overnight at 42°C with the 5′-end-labeled ^32^P-TelG probe in Church buffer, washed twice for 5 min each with 0.2 M wash buffer (0.2 M Na_2_HPO4 pH 7.2, 1 mM EDTA, and 2% SDS) at room temperature, and once for 10 min with 0.1 M wash buffer at 42°C. Signals were analyzed using a phosphor-imager, visualized on a Typhoon 9410 Imager (GE Healthcare), and processed with ImageQuant 5.2 software (Molecular Dynamics).

For detection with the DIG-TelG probe, hybridization and chemiluminescent detection was performed as previously described^64^. Briefly, membranes were pre-hybridized for 1 hour at 42°C in DIG Easy Hyb^TM^ (11603558001, Roche) and then hybridized overnight with the DIG-TelG probe. After hybridization, membranes underwent stringent washes - twice with 2x SSC containing 0.1% SDS and twice with 0.5x SSC with 0.1% SDS - at 42°C. For signal detection, membranes were blocked with 1% blocking reagent (11096176001, Roche) in maleic acid buffer (pH 7.5) and incubated with an anti-digoxigenin-peroxidase (POD) conjugated antibody (1:1,000, 11207733910, Roche). Telomeric signals were detected using a peroxidase-mediated chemiluminescence substrate (Cytiva). High-resolution digital images were captured using the Azure Biosystems Imaging System.

### Short Multiply Aggregated Sequence Homologies (SMASH)

Copy number profiles for U2OS cell line were generated as previously described^37^. Briefly, genomic DNA was fragmented using dsDNA Fragmentase (M0348, NEB) to produce fragments with an average size of approximately 40 bp. The resulting short fragments were randomly ligated to generate chimeric DNA molecules ranging from ∼400–700 bp. Fragments within the desired size range were purified using Agencourt AMPure XP beads (A63881, Beckman Coulter) and subsequently processed for Illumina library preparation using NEBNext Multiplex Dual Index Adaptors and Primer Sets (E6440, NEB). Libraries were sequenced on an Illumina MiSeq platform. Sequencing reads were demultiplexed and aligned to the reference genome to generate copy number and karyotype profiles as previously described^37^.

### Clonogenic-based growth assay

To evaluate difference in growth capacity, indicated control and experimental cells were plated at densities of 10,000 - 25,000 cells per well of 6-well plates, or 2,500 cells per well in 24-well plates. Cells were cultured with their respective growth media in a humidified incubator at 37°C with 5% CO2 for 10 - 20 days, after which they were stained with 0.5% crystal violet in 20% methanol for 30 minutes.

### Metaphase spread preparation and telomere FISH

Cells synchronized in G2/M phase by 10 μM CDK1 inhibitor Ro-3306 (S7747, Selleckchem) were washed with warmed PBS and released into fresh medium (pre-warmed to 37 °C) containing 0.1 μg/mL Karyomax/Colcemid (Life Technologies) for 90 min. Mitotic cells were then harvested, incubated with 75 mM KCl (pre-warmed to 37°C) for 25 min at 37°C, and prefixed with several drops of ice-cold fixation buffer (methanol:acetic acid, 3:1). Cells were pelleted, resuspended in ice-cold fixation buffer, and fixed for 30 min. After three washes with fixation buffer, cells were dropped onto ethanol-water-cleaned slides. Following air-drying overnight, telomere signals were detected by FISH. Images were captured using a 60x lens on a Nikon Eclipse Ti2-E live-cell and Deconvolution Imaging System.

### RNA extraction and quantitative RT-PCR (RT-qPCR)

Total RNA was extracted from cells using the NucleoSpin RNA kit (Macherey-Nagel). Reverse transcription was carried out on 1.0 μg of total RNA using the RevertAid RT kit (Thermo Fisher Scientific). RT-qPCR was performed on cDNA samples using PowerUp SYBR Green Master Mix (Thermo Fisher Scientific) on a 7500 Fast Real-time PCR system (Thermo Fisher Scientific). All samples were run in triplicate, and mRNA levels were normalized to GAPDH mRNA. Relative mRNA expression levels were presented as 2^ΔΔCt^ values.

### RNA-seq and pathway analysis

Total RNA was extracted using the NucleoSpin RNA isolation kit, and subsequent sequencing was conducted at the Weill Cornell Medicine Genomics Core facility. Libraries were constructed using the Illumina TruSeq stranded mRNA kit. Raw paired-end reads were processed for quality control and adapter removal using bbduk with the following parameters: ktrim=r, k=23, mink=11, hdist=1, qtrim=rl, trimq=10, maq=10. Read quality was assessed using FastQC (v0.12.1). Processed reads were mapped to the human reference genome (GRCh38/hg38) using STAR (v2.7.11b) in 2-pass Basic mode. Gene-level quantification for reverse-stranded exons was performed with featureCounts (Subread v2.0.6; settings: -p -t exon -g gene_id -s 2). Following BAM indexing via samtools (v1.19.2), differential expression analysis was executed using DESeq2 (v1.44.0). To account for inter-clonal experimental variability (biological replicates), a design formula of ∼ batch + class was applied. Genes with low abundance (< 10 counts in fewer than 2 samples) were excluded. Dispersion was estimated and models were fitted using default DESeq2 parameters with independent filtering. For downstream computational analyses, variance-stabilizing transformation (blind = FALSE) was utilized.

Genes were ranked by the ratio between comparison samples and subjected to functional pathway enrichment analysis using GSEA (GSEA v4.4.0, Broad Institute)^65^. Regular GSEA was performed on publicly available datasets GSE141331 and GSE141335. All analyses were run with 1,000 permutations, and gene sets with nominal p < 0.05 and FDR q < 0.1 were considered significantly enriched.

### Chromatin Immunoprecipitation (ChIP)

ChIP assays were performed as described previously^36^. In brief, cells (up to approximately 2 × 10^7^) were cross-linked with 1% formaldehyde for 10 minutes at room temperature, followed by quenching with 125 mM glycine. Chromatin DNA was fragmented to an average size of 200–500 bp using a focused ultrasonicator M2 (Covaris). For immunoprecipitation, solubilized chromatin was incubated overnight at 4°C with 5 μg of anti-STAG2 antibody (#4464, abcam) or normal goat IgG as a control. Immune complexes were captured using Protein G magnetic beads, washed with RIPA buffer, and eluted. Cross-links were reversed at 65°C for 15 hours, and the resulting DNA was purified using a PCR purification kit (MN) for downstream telomeric analysis.

### Telomere Dot Blot Analysis (DIG-System)

To quantify telomeric DNA associated with STAG2, a non-radioactive dot blot assay was performed using the digoxigenin (DIG) system. Purified ChIP DNA and corresponding input DNA were diluted in 0.4N NaOH containing 10 mM EDTA, denatured at 95°C for 10 min, and loaded onto a nylon membrane using a vacuum dot-blot apparatus. DNA was fixed to the membrane by UV cross-linking at 125 mJ.

Membranes were pre-hybridized for 1 hour at 42°C in DIG Easy HybTM (11603558001, Roche) and then hybridized overnight with a DIG-labeled TelG oligonucleotide probe. Following hybridization, membranes underwent stringent washes - twice 2x SSC with 0.1% SDS and twice 0.5x SSC with 0.1% SDS - at 42°C.

For signal detection, membranes were blocked with 1% blocking reagent (11096176001, Roche) in maleic acid buffer (pH 7.5) and incubated with an anti-digoxigenin-peroxidase (POD) conjugated antibody (1:1,000; 11207733910, Roche). Telomeric signals were detected using a peroxidase-mediated chemiluminescence substrate (Cytiva). High-resolution digital images were captured using the Azure Biosystems Imaging System.

### Protein expression and purification

cDNAs encoding human STAG2 (Uniprot: Q8N3U4), PAGR1 (Uniprot: Q9BTK6), RAD21 (Uniprot: O60216), and PAXIP1 (Uniprot: Q6ZW49) were synthesized and codon-optimized for expression in *Spodoptera frugiperda* (Sf9) cells. STAG2 was cloned into the pFastBac1 vector with a C-terminal 8xHis and Twin-StrepII dual tag. PAGR1 was cloned into a vector with an N-terminal 6xHis tag. RAD21 (residues 281-420aa) was cloned into the pFastBac1 vector with an N-terminal 10xHis and Twin-StrepII dual tag. PAXIP1 (residues 94-183aa) cloned into the pFastBac1 vector with an N-terminal 10xHis and Twin-StrepII dual tag.

Protein expression was used with the Bac-to-Bac™ Baculovirus Expression System (Thermo Fisher). Briefly, the constructed plasmids were transformed into DH10Bac™ *E. coli*. High-molecular-weight recombinant bacmid DNAs were isolated and transfected into Sf9 cells using PEI Reagent (23966-1, Kyfora Bio) to generate recombinant baculoviruses suitable for preliminary expression experiments.

For co-expression, STAG2, PAGR1, RAD21 (281-420aa), and PAXIP1 (94-183aa) were expressed together in High Five cells (Thermo Fisher). Cells were infected with recombinant baculovirus at a density of 2.0×10^6^ cells/ml and incubated at 27°C with shaking at 120 rpm for 72 hours. At a cell viability of 70-80%, the cells were harvested and stored at −80°C until use. Pellets from 900 ml High Five cells were resuspended in lysis buffer (50 mM HEPES-KOH pH 8.0, 500 mM KCl, 0.5 mM EDTA, 0.5 mM TCEP, 5% (w/v) glycerol, Protease Inhibitor). Cells were lysed by sonication for 8 min and centrifuged at 48,000 g for 30 min at 4°C. The supernatant was incubated with 2 ml Strep-Tactin®XT 4Flow® resin (IBA, 2-5010-025) for 30 min at 4°C. The resin was first washed with lysis buffer, followed by lysis buffer containing 1.2 M KCl. Proteins were eluted with lysis buffer containing 50 mM Biotin. The eluate was concentrated and fractionated on a Superdex 200 Increase 10/300 GL column (28990944, Cytiva) pre-equilibrated with buffer (20 mM HEPES, pH 8.0, 300 mM KCl, 0.5 mM TCEP). The purified complexes were collected and prepared for the cryo-EM analysis.

### Cryo-EM grid preparation and data acquisition

The purified STAG2/SCC1/PAGR1/PAXIP1 complex sample at a concentration of approximately 1 mg/ml was applied (3 μl) to a glow-discharged Quantifoil R1.2/1.3 300-mesh gold holey carbon grid. The grid was blotted for 5 seconds with a blot force of 4 under 100% humidity at 8°C and plunged into liquid ethane using a Vitrobot Mark IV (Thermo Fisher). The grid was initially screened using a 200-kV cryo-electron microscope before high-resolution data collection. Micrographs were acquired on a 300-kV cryo-electron microscope (Titan Krios G4, Thermo Fisher Scientific) equipped with a Gatan K3 directed electron detector and a GIF BioQuantum energy filter. Data were collected in super-resolution mode with 40 frames per movie over 3 seconds, at a total accumulated dose of 50 electrons per Å^2^. All images were acquired in counting mode at a nominal magnification of 105,000 ×, corresponding to a calibrated physical pixel size of 0.824 Å. The defocus range was set between - 1.0 to −3.0 μm. Detailed cryo-EM data collection parameters are provided in Extended Data Table 1.

### Cryo-EM structural determination

Cryo-EM datasets were processed with CryoSPARC v4.6.2^66^. Movie frames were pre-processed by CryoSPARC live. The Patch Motion Correction and the Patch CTF estimation program were used for motion correction and contrast transfer function estimation. 4263 micrographs of the PAXIP1-PAGR1-STAG2-RAD21 complex were exported from CryoSPARC live work session into CryoSPARC workspace. 3171,200 particles were automatically picked by Blob Picker and inspect picks. Multiple rounds of 2D classification for initial particle picking. A total of 1324,021 particles were classified into three classes after ab-initio reconstruction and heterogeneous refinement. The best class of 785,386 particles was carried out and extracted using a box size of 320 pixels for non-uniform refinement, which generated a 2.51 Å map based on the Fourier shell correlation (FSC). By using a mask focused on the interaction between STAG2 and PAGR1, we improved the local density and obtained a map at 2.49 Å resolution.

### Model building and refinement

The structure of the PAXIP1-PAGR1-STAG2-RAD21 complex was predicted using AlphFold3^67^, and the resulting model was used as an initial template. The model was docked into the cryo-EM map using ChemiraX^68^, manually adjusted in Coot^69^, and refined in Phenix^70^. Real-space refinement in Phenix was used to perform the refinement for all the structural data (Extended Data Table 1). All structural figures were generated by ChemiraX.

### Biolayer interferometry (BLI) binding assay

BLI assays were performed on a ForteBio Octet RED96 (Sartorius). Streptavidin (SA) sensors were first equilibrated in BLI assay buffer (Buffer BLI) containing 20 mM HEPES, pH 8.0, 150 mM KCl, and 0.02% Tween20. SA sensors were loaded with STAG2-RAD21 for 5 min at a 4 μg/ml concentration, which was labeled with NHS-PEG12-Biotin (A35389, Thermo Fisher Scientific) according to the manufacturer’s instructions. The biotin-labeled STAG2 sensors were dipped into different concentrations (0.25-8 μM) of PAGR1 for association, followed by dipping into assay buffer for dissociation. The control sensor was a biotin-labeled STAG2 sensor associated with the BLI assay buffer only. The data was analyzed with the Octet systems software.

### Statistics and Reproducibility

Details regarding quantitation and statistical analysis are provided in the figures and figure corresponding legends. All statistical analysis were performed using GraphPad Prism software (version 7.0e). Significance was calculated using an unpaired two-tailed Student’s t-test. P values are shown in the figures.

Sample sizes were determined based on preliminary data; no statistical methods were used to predetermine sample sizes. All results are presented as mean ± s.e.m. Normality of sample distribution was verified before applying the two-tailed t test for two group comparisons. Differences were considered significant when P < 0.05.

For western blotting, each experiment was repeated independently at least twice with similar results; representative images/blots are shown. The experiments were not randomized, and the investigators were not blinded to allocation during experiments or outcome assessment.

## Data availability

The data supporting the findings of this study are available within the article and its Supplemental Tables. RNA-seq data have been deposited in the Gene Expression Omnibus (GEO) database under accession number GSE314012; SMASH data are available in the NCBI Sequence Read Archive (SRA; http://www.ncbi.nlm.nih.gov/sra/) under accession number SRX32945314. Cryo-EM density and atomic coordinates for the PAXIP1-PAGR1-STAG2-RAD21 complex have been deposited in the Protein Data Bank under the access codes pdb_000043lt (PDB ID 43LT) and EMDB entry ID EMD-81970.

## Code availability

For specific requests, please contact the corresponding authors.

## Acknowledgements

We thank A. Marzio (Weill Cornell) for technical assistance with the Nikon Ti2-E imaging system. This work was partially supported by an Andrew McDonough B+ Foundation research award to J.P., and the William Rhodes and Louise Tilzer-Rhodes Center for Glioblastoma Research Award to H.Z., J.P., and N.F.L. Z.W. and the SMASH component of this study was supported by the Simons Foundation (SFARI Award No. 17457).

## Author contributions

J.P., C.D., and H.Z. designed the experiments. S.Y.H. and X.H. performed the majority of cellular experiments, with assistance from I.M., S.K., S.Y., Y.K., and Y.W. X.W. carried out cryo-EM sample preparation and structure determination, with help from Y.X. and Z.Z. F.L. initiated the project and constructed the conditional DAXX knockout system. Z.W. performed SMASH assay and data analysis, with assistance from B.Y. E.Y.Y. performed TRF analysis under the supervision of N.F.L. J.P. analyzed the RNA-seq data. J.P., C.D., and H.Z. wrote the manuscript with input from all other authors.

---

**Extended Data Fig. 1.**
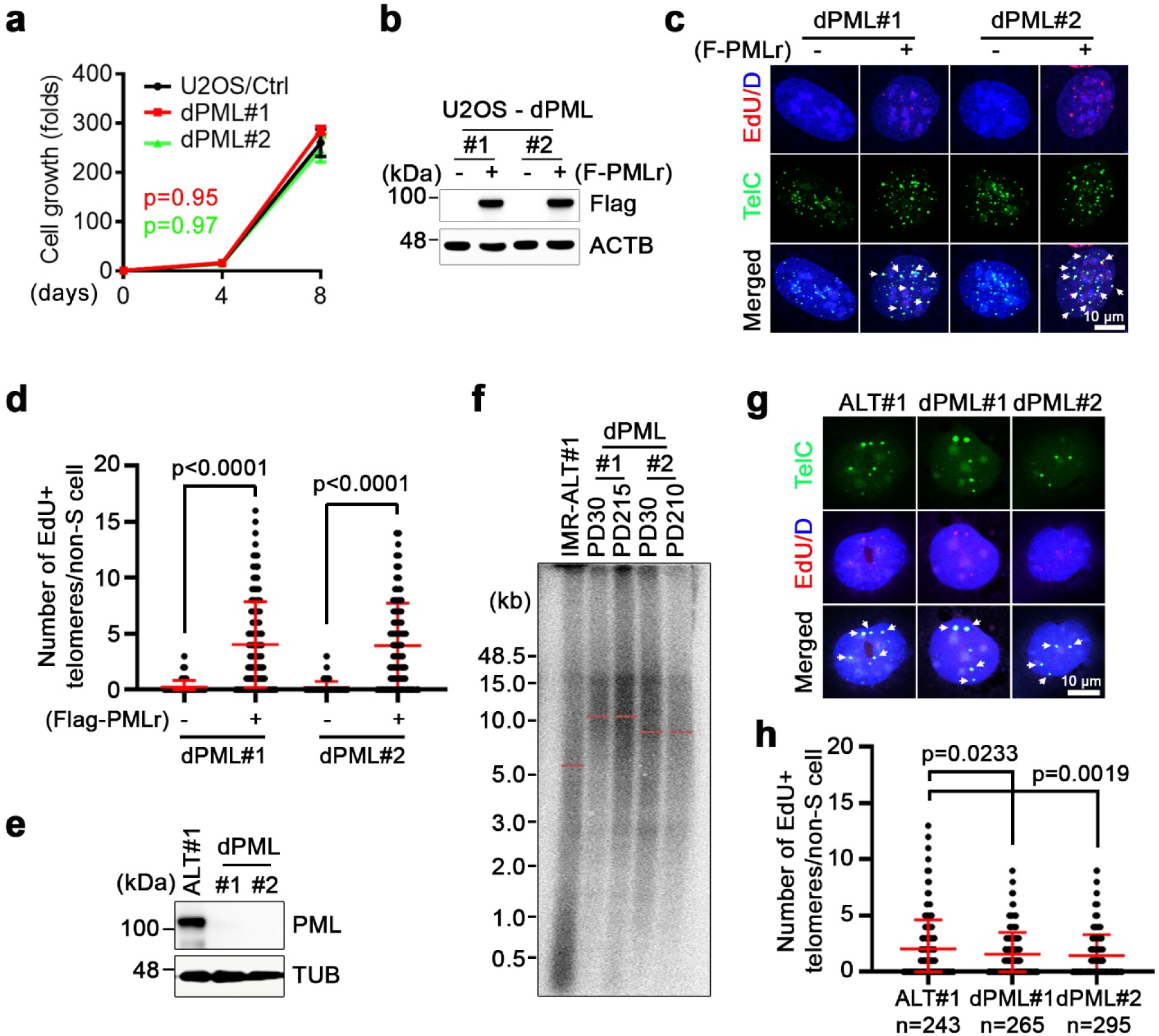
PML is dispensable for ALT-mediated telomere length maintenance. **a,** Cell proliferation assays of parental U2OS control cells and two CRISPR-derived PML deletion clones (dPML#1 and #2). Data represent mean ± s.e.m. from three independent experiments. Statistical analysis was performed using an unpaired two-tailed Student’s t test; P values are shown. **b,** Western blot analysis of ectopically expressed, CRISPR-resistant, and Flag-tagged PML (Flag-PMLr, detected with anti-Flag antibody) in U2OS-dPML clones (#1 and #2) transduced with either vector control (−) or Flag-PMLr (+). ACTB serves as a loading control. **c,** Representative immunofluorescence (IF)-FISH images showing non-S-phase EdU colocalization with telomeres (TelG) in U2OS-dPML clones (#1 and #2) transduced with either vector control (−) or Flag-PMLr (+). Arrows denote EdU-positive telomere foci. **d,** Quantification of (**c**), showing the number of non-S-phase EdU-positive telomere foci per cell in U2OS-dPML clones (#1 and #2) transduced with either vector control (−) or Flag-PMLr (+). Data represent mean ± s.e.m. from three independent experiments. Statistical analysis was performed using an unpaired two-tailed Student’s t test; P values are shown. **e,** Western blot analysis of PML protein expression in parental IMR90-ALT#1 cells and two CRISPR/Cas9-derived PML deletion clones (dPML#1 and #2). **f,** TRF analysis of telomere length in parental IMR90-ALT#1 cells and dPML clones (#1 and #2) at the indicated population doublings (PD). Genomic DNA prepared from the indicated cells were assayed by a ^32^P-labeled TelG probe. **g,** Representative IF-FISH images showing non-S-phase EdU colocalization with telomeres (TelG) in parental IMR90-ALT#1 cells and dPML clones (#1 and #2). Arrows denote EdU-positive telomere foci. **h,** Quantification of (**g**), showing the number of non-S-phase EdU-positive telomere foci per cell in parental IMR90-ALT#1 cells and dPML clones. Data represent mean ± s.e.m. from three independent experiments. Statistical analysis was performed using an unpaired two-tailed Student’s t test; P values are shown.

**Extended Data Fig. 2.**
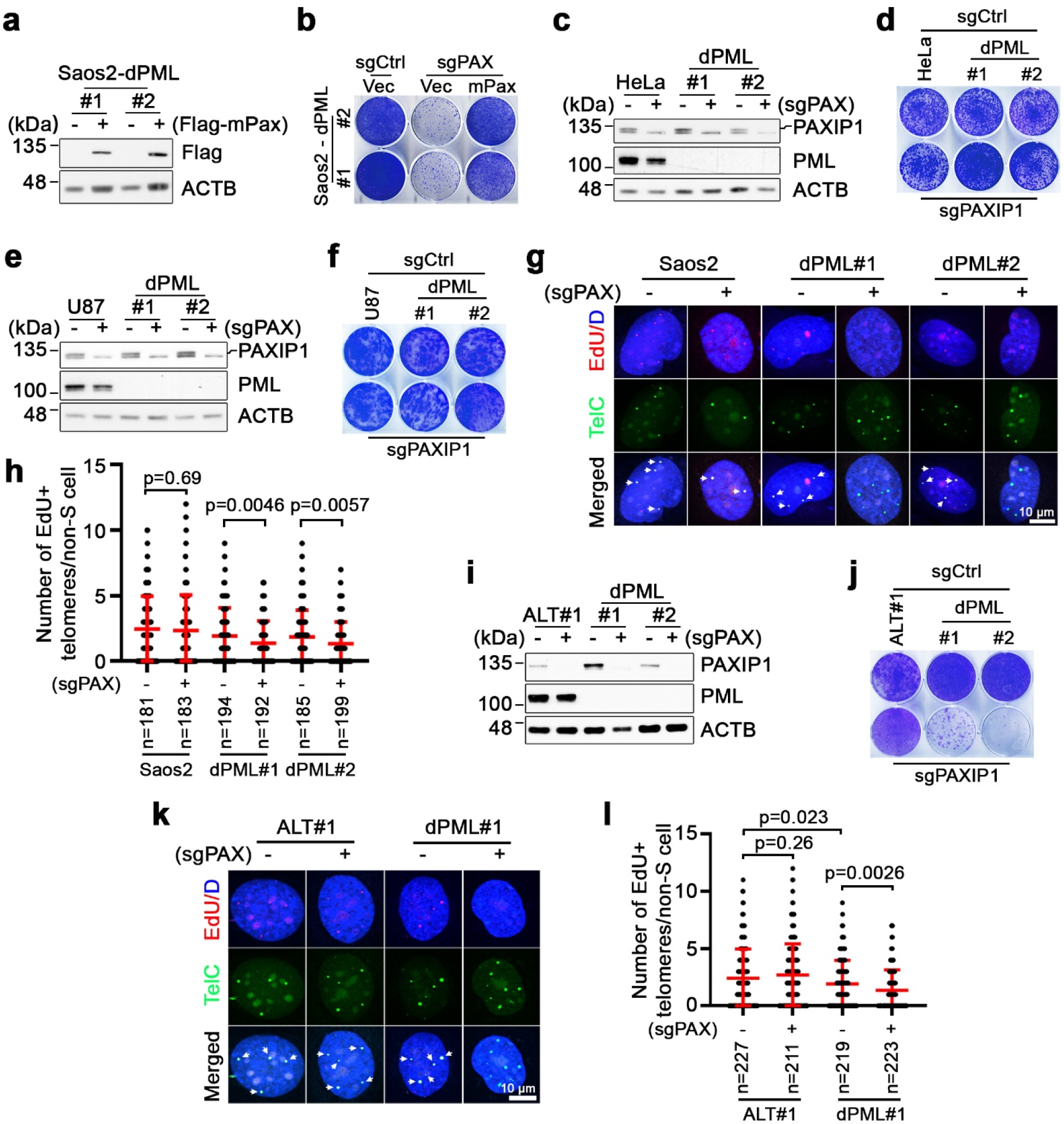
Synthetic lethal interaction between PAXIP1-PAGR1 and PML occurs specifically in ALT-positive cells. **a,** Western blot analysis of ectopically expressed Flag-tagged murine Paxip1 (Flag-mPax) in Saos2-dPML clones (#1 and #2) transduced with either vector control (−) or Flag-mPax (+). **b,** Crystal violet-based clonogenic assay of Saos2-dPML clones (#1 and #2) complemented with either vector (Vec) or Flag-mPax (mPax) and further transduced with sgCtrl or sgPAXIP1 (sgPAX). Crystal violet staining was performed on day 18 post-seeding. **c,** Western blot analysis of PAXIP1 and PML protein expression in parental HeLa cells and CRISPR/Cas9-derived PML deletion clones (dPML#1 and #2) following transduction of sgCtrl (−) or sgPAXIP1 (+). **d,** Crystal violet-based clonogenic assay of parental HeLa cells and HeLa-dPML#1 and #2 cells transduced with sgCtrl (−) or sgPAXIP1 (+). **e,** Western blot analysis of PAXIP1 and PML protein expression in parental U87 cells and CRISPR/Cas9-derived PML deletion clones (dPML#1 and #2) following transduction of sgCtrl (−) or sgPAXIP1 (+). **f,** Crystal violet-based clonogenic assay (F) of parental U87 cells and U87-dPML#1 and #2 cells transduced with sgCtrl (−) or sgPAXIP1 (+). **g,** Representative IF-FISH images of non-S-phase EdU colocalization with telomeres (TelG) in parental Saos2 cells and Saos2-dPML clones transduced with sgCtrl or sgPAX. Arrows denote EdU-positive telomere foci. **h,** Quantification of (**g**), showing the number of non-S-phase EdU-positive telomere foci per cell in parental Saos2 cells and Saos2-dPML clones transduced with sgCtrl or sgPAX. Data represent mean ± s.e.m. from three independent experiments. Statistical analysis was performed using an unpaired two-tailed Student’s t test; P values are shown. **i-k,** Western blot analysis of PAXIP1 and PML protein expression (**i**), crystal violet-based clonogenic assay (**j**), and representative IF-FISH images showing non-S-phase EdU colocalization with telomeres (**k**) from parental IMR90-ALT#1 cells and dPML clones following transduction with sgCtrl (−) or sgPAXIP1 (sgPAX). **l,** Quantification of (**k**), showing the number of non-S-phase EdU/telomere colocalization foci per cell in parental IMR90-ALT#1 cells and the dPML#1 clone transduced with sgCtrl or sgPAXIP1. Data represent mean ± s.e.m. from three independent experiments. Statistical analysis was performed using an unpaired two-tailed Student’s t test; P values are shown.

**Extended Data Fig. 3.**
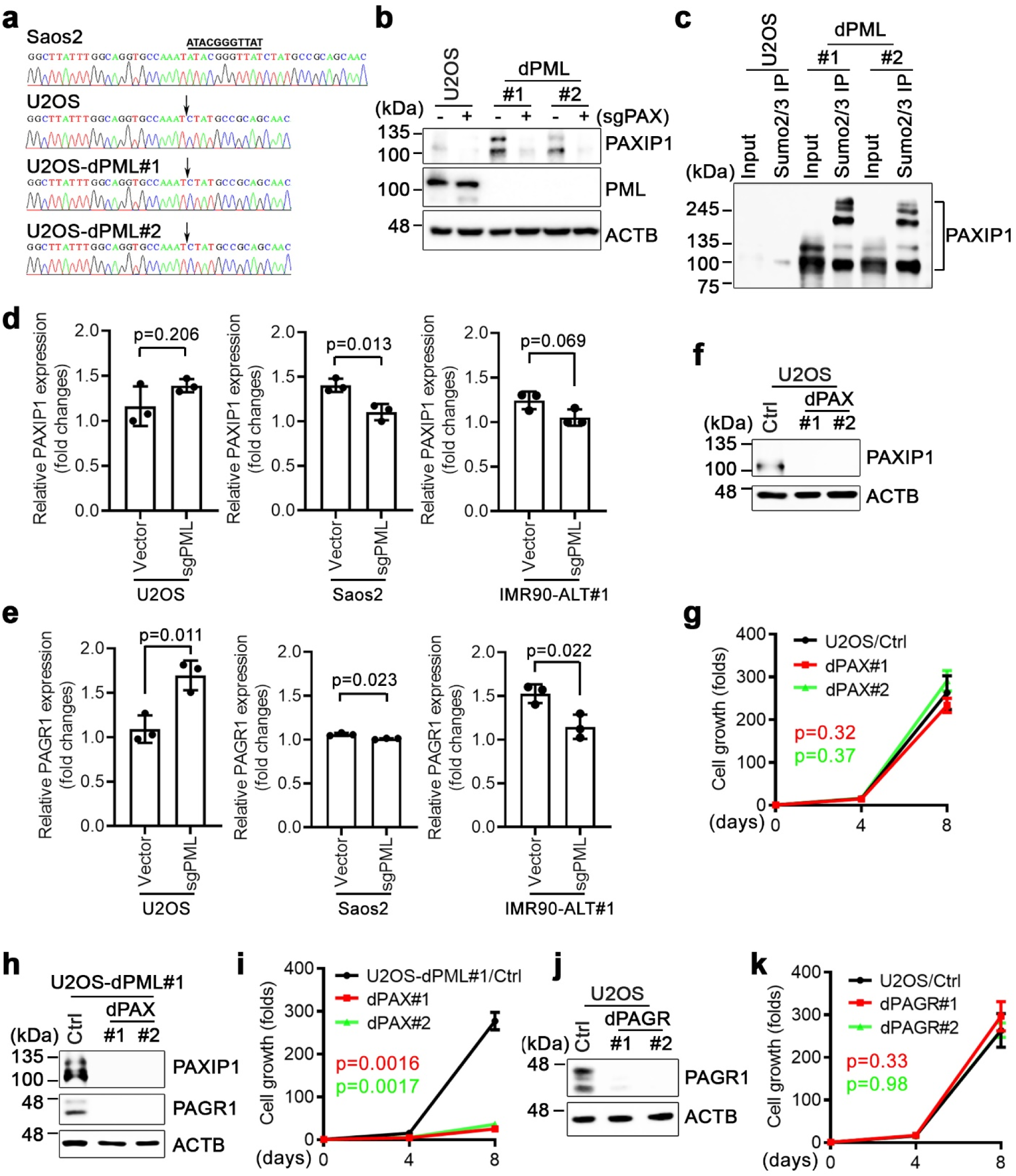
PAXIP1-PAGR1 and PML cooperate to promote ALT telomere maintenance. **a,** Sanger sequencing traces showing wild type PAXIP1 sequence in Saos2 cells and the PAXIP1 deletion (c.2204_2214) in parental U2OS cells and U2OS-dPML clones after 450 population doublings in culture. The indel is indicated. **b,** Western blot analysis of PAXIP1 and PML protein expression in parental U2OS cells and dPML clones (#1 and #2) following transduction with sgCtrl (−) or sgPAXIP1 (+). **c,** Western blot analysis of input and SUMOylated PAXIP1 protein levels in parental U2OS cells and dPML clones (#1 and #2). SUMOylated protein were captured by immunoprecipitation (IP) using an anti-SUMO2/3 antibody under a denaturing condition. **d,** RT-qPCR analysis of *PAXIP1* mRNA expression in U2OS, Saos2, or IMR90-ALT#1 cells following transduction with sgCtrl or sgPML. **e,** RT-qPCR analysis of *PAGR1* mRNA expression in U2OS, Saos2, or IMR90-ALT#1 cells following transduction with sgCtrl or sgPML. **f,** Western blot analysis of PAXIP1 protein expression in parental U2OS cells and CRISPR/Cas9-derived PAXIP1 deletion clones (dPAX#1 and #2). **g, C**ell proliferation assays of parental U2OS cells (Ctrl) and U2OS-dPAX (#1 and #2) cells. Data represent mean ± s.e.m. from three independent experiments. Statistical analysis was performed using an unpaired two-tailed Student’s t test; P values are shown. **h,** Western blot analysis of PAXIP1 and PAGR1 protein expression in U2OS-dPML#1 cells and CRISPR/Cas9-derived PAXIP1 deletion clones (dPAX#1 and #2). **i,** Cell proliferation assays of U2OS-dPML#1 cells (Ctrl) and dPAX (#1 and #2) cells. Data represent mean ± s.e.m. from three independent experiments. Statistical analysis was performed using an unpaired two-tailed Student’s t test; P values are shown. **j,** Western blot analysis of PAGR1 protein expression in parental U2OS cells and CRISPR/Cas9-derived PAGR1 deletion clones (dPAGR#1 and #2). **k,** Cell proliferation assays of parental U2OS cells (Ctrl) and dPAGR (#1 and #2) cells. Data represent mean ± s.e.m. from three independent experiments. Statistical analysis was performed using an unpaired two-tailed Student’s t test; P values are shown.

**Extended Data Fig. 4.**
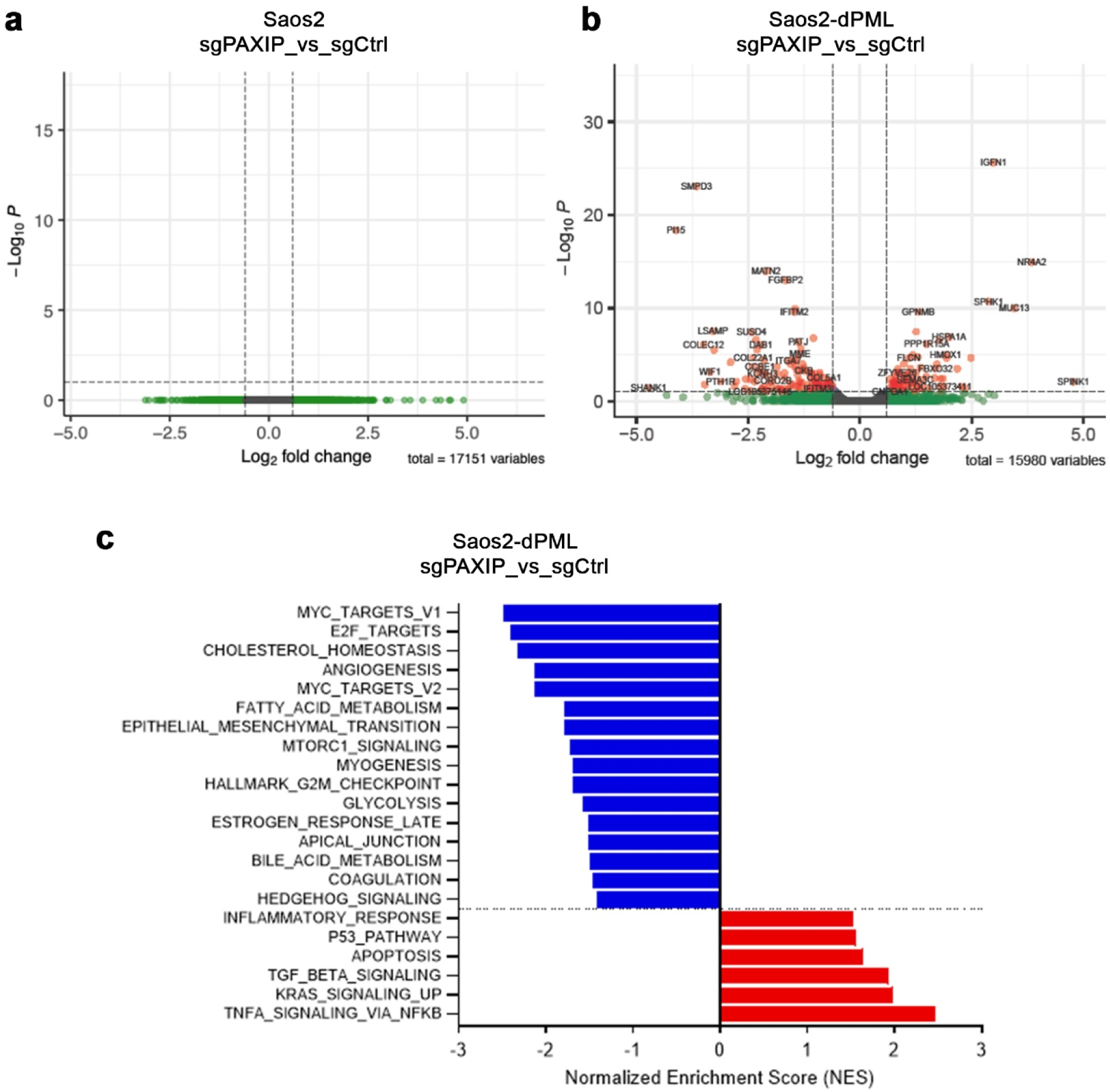
PAXIP1 depletion has minor impact on the transcription of genes associated with DNA damage repair. **a, b,** Volcano plots of RNA-Seq data comparing sgPAXIP1 versus sgCtrl in Saos2 cells (**a**) or in Saos2-dPML cells (**b**). Negative log2 fold change indicates lower expression in sgPAXIP1-transduced cells. Genes with significantly changed mRNA expression (-log_10_ FDR > 2) are shown as red dots. **c,** Top significant enrichments identified by gene set enrichment analysis (GSEA) from the 224 differentially expressed genes comparing Saos2-dPML cells transduced with sgPAXIP1 versus sgCtrl.

**Extended Data Fig. 5.**
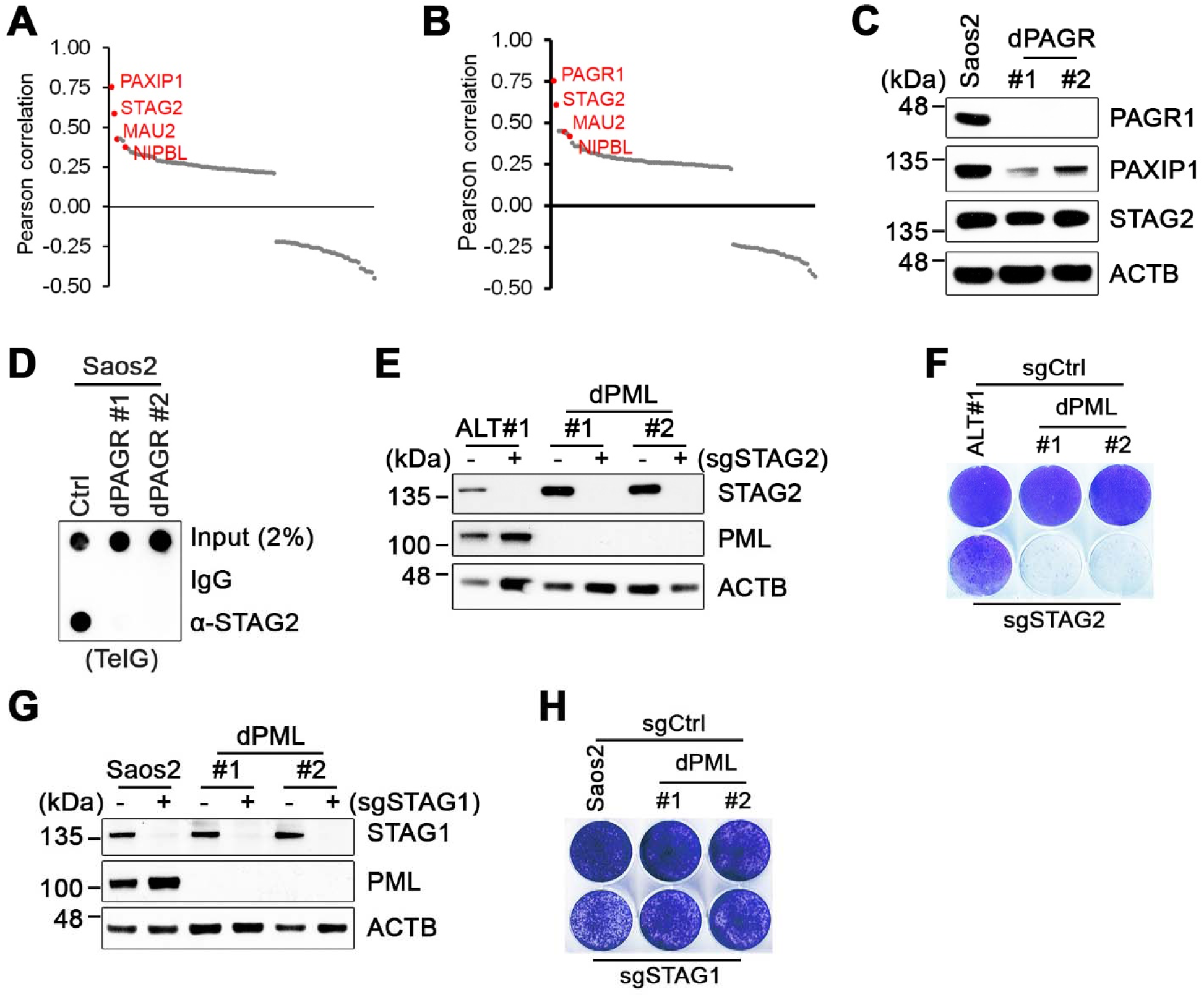
STAG2 functions epistatically with the PAXIP1-PAGR1 complex in ALT-mediated telomere maintenance. **a, b,** Pearson correlation coefficients of gene dependency scores for the top 100 co-dependencies of PAXIP1 (**a**) and PAGR1 (**b**). Scores are based on the CRISPR Avana database from the DepMap portal (https://depmap.org). **c, d,** Western blot analysis of PAGR1, PAXIP1, and STAG2 protein expression (**c**) and telomere dot-blot analysis of anti-STAG2 ChIP (**d**) in parental Saos2 cells and two CRISPR/Cas9-derived PAGR1 deletion clones (dPAGR#1 and #2). **e, f,** Western blot analysis of STAG2 and PML protein expression (**e**) and crystal violet-based clonogenic assay (**f**) of parental IMR90-ALT#1 cells and dPML clones (#1 and #2) following transduction with sgCtrl (−) or sgSTAG2 (+). **g, h,** Western blot analysis of STAG1 and PML protein expression (**g**) and crystal violet-based clonogenic assay (**h**) of parental Saos2 cells and dPML clones (#1 and #2) following transduction with sgCtrl (−) or sgSTAG1 (+).

**Extended Data Fig. 6.**
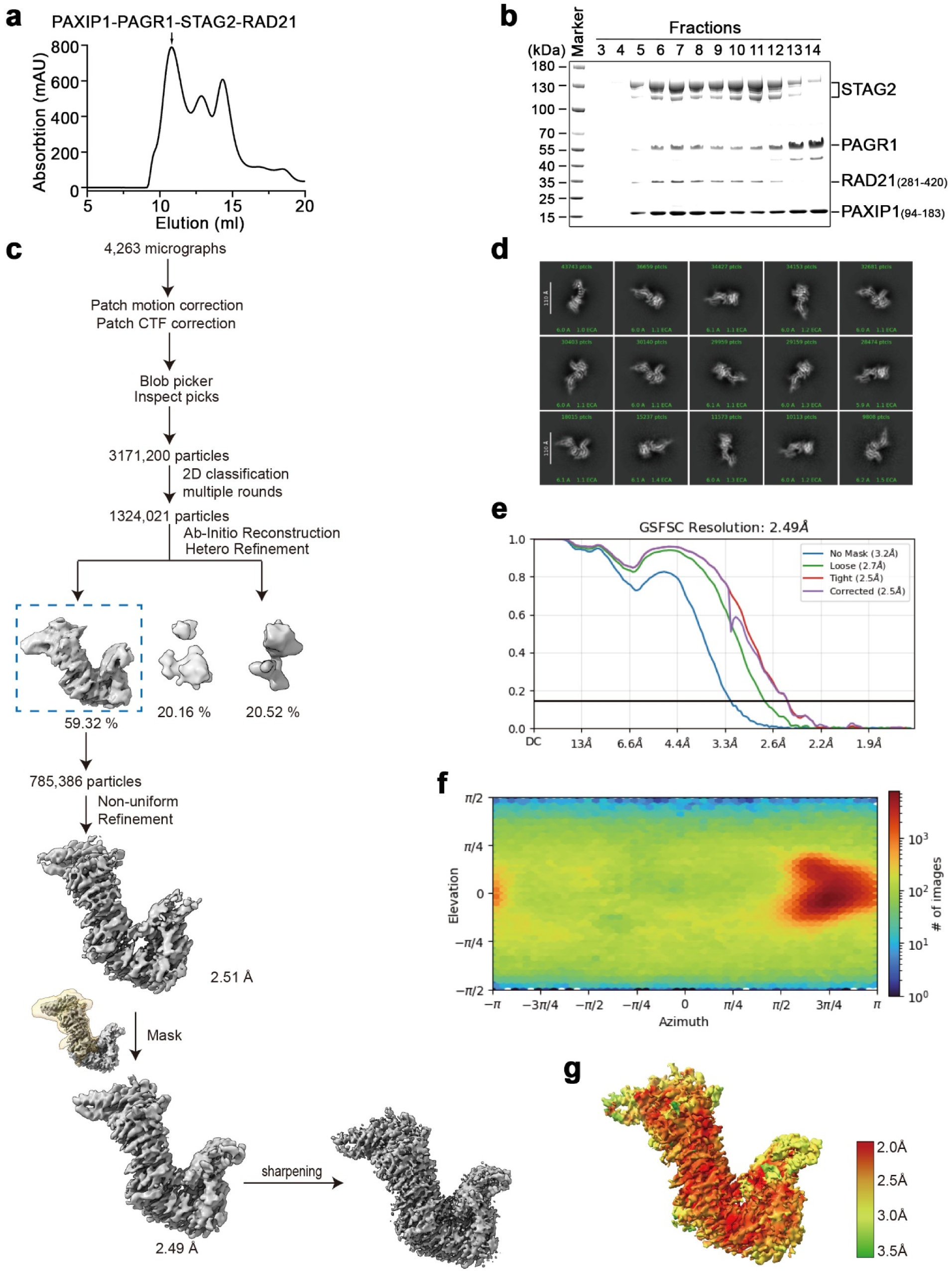
Cryo-EM structure determination of the human PAXIP1-PAGR1-STAG2-RAD21 complex. **a,** Size exclusion chromatography elution profile of the human PAXIP^194–183^-PAGR1-STAG2-RAD21^281–420^ complex on a Superdex 200 Increase 10/300 GL column. The peak indicated by the black arrow corresponds to the fraction used for cryo-EM sample preparation. **b,** Coomassie blue-stained SDS-PAGE gel of the PAXIP^194–183^-PAGR1-STAG2-RAD21^281–420^ complex purified by gel filtration. **c,** Cryo-EM processing pipeline for the PAXIP1-PAGR1-STAG2-RAD21 complex. Approximately 3 million particles were selected from 4,263 micrographs and cleaned up by multiple rounds of 2D classification. The final set of 785,386 particles were subjected to non-uniform refinement, generating a 2.51Å map. Local refinement was performed using a mask around the STAG2-PAGR1 interface, yielding a map at 2.49Å resolution. **d,** 2D class averages of the PAXIP1-PAGR1-STAG2-RAD21 complex. Scale bars, 100 Å. **e,** Gold-standard Fourier shell correlation (FSC) curve of the PAXIP1-PAGR1-STAG2-RAD21 complex. **f,** Angular distribution plot of the PAXIP1-PAGR1-STAG2-RAD21 complex. **g,** Cryo-EM density maps of the PAXIP1-PAGR1-STAG2-RAD21 complex, color-coded according to local resolution ranging from 2.0 Å to 3.5 Å.

**Extended Data Fig. 7.**
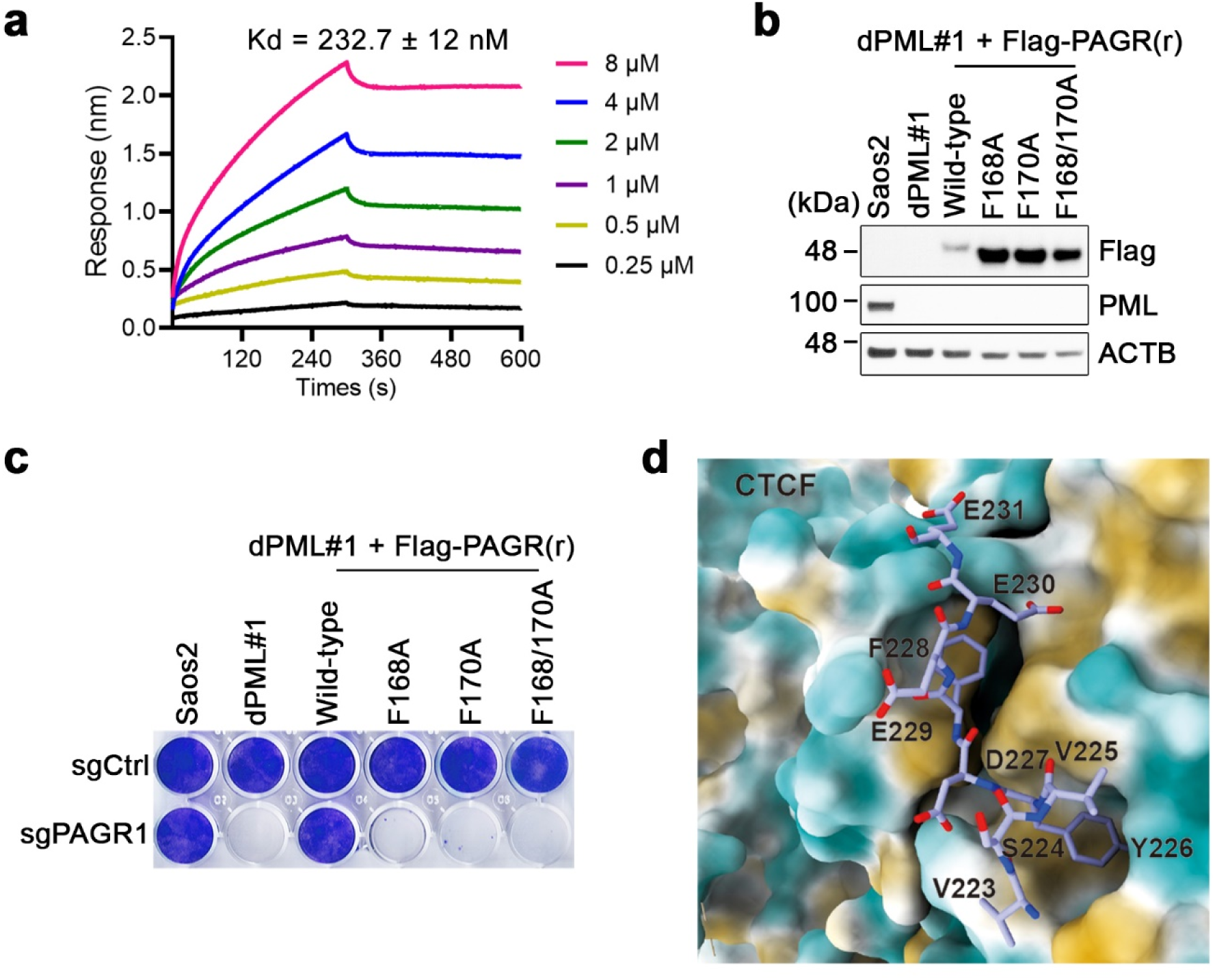
PAGR1 FDF motif is critical for its ALT-directed telomere maintenance function. **a,** Binding kinetics of PAGR1 with STAG2-RAD21 (281-420) cohesin subcomplex determined by biolayer interferometry (BLI) assays. **b,** Western blot analysis of ectopically expressed CRISPR-resistant Flag-tagged PAGR1 (wild-type or mutant, detected with anti-Flag antibody) and along with PML protein expression in parental Saos2 cells and the dPML#1 clones transduced with the indicated constructs. ACTB serves as a loading control. **c,** Crystal violet-based clonogenic assay of parental Saos2 cells and dPML clones expressing the indicated construct following further transduction with either sgCtrl or sgPAGR1. Crystal violet staining was performed on day 18 post-seeding. **d,** Hydrophobic pockets of STAG2-RAD21 cohesin subcomplex interact with CTCF. Residues Y226 and F228 of CTCF bind to the respective hydrophobic pockets formed jointly by STAG2 and RAD21.

**Extended Data Fig. 8.**
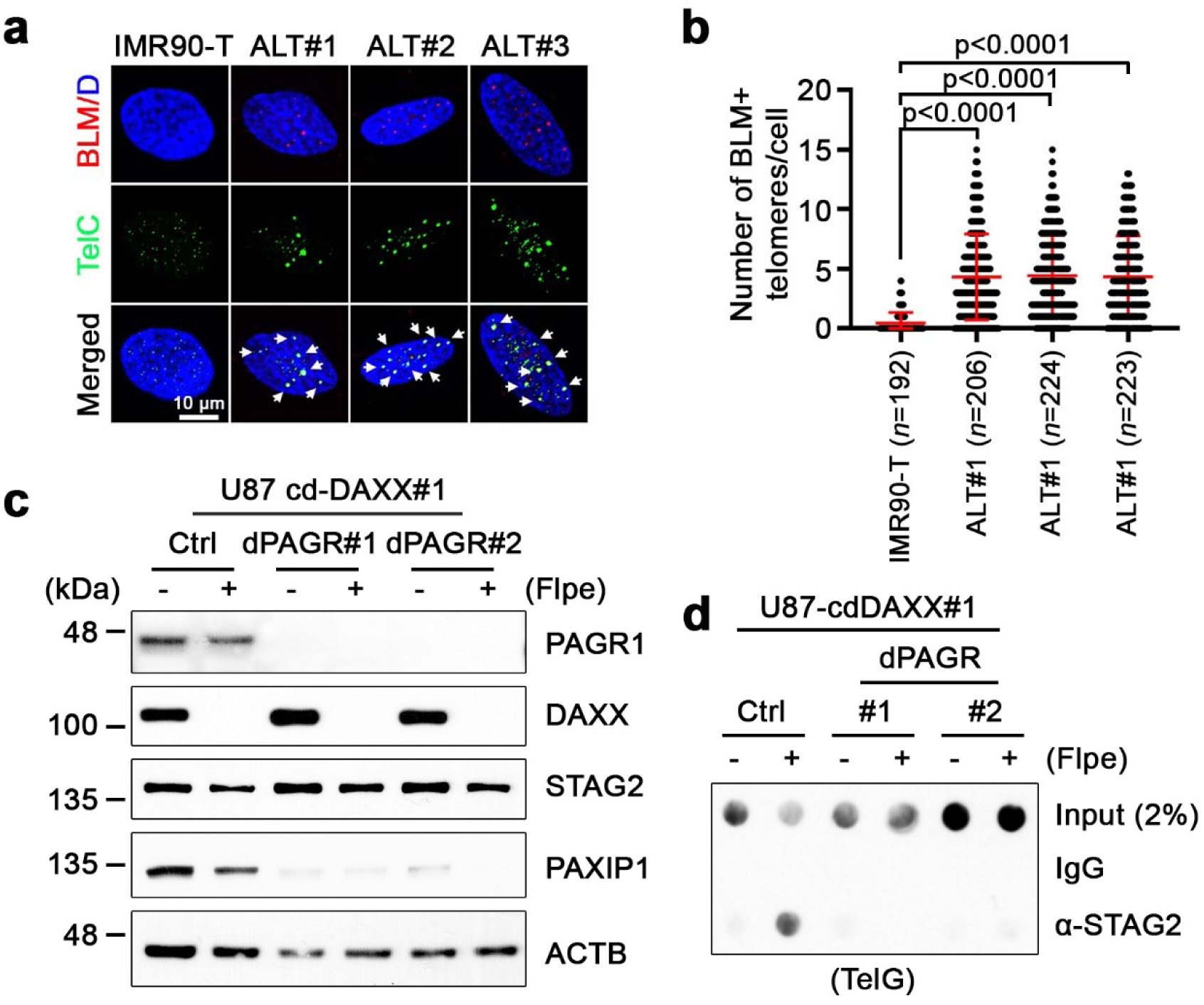
PAGR1 is required for *de novo* telomere cohesin recruitment during break-induced telomere repair. **a,** Representative IF-FISH images showing BLM colocalization with telomeres (TelG) in G2/M phase-synchronized IMR90-T and IMR90-ALT cells. Arrows denote BLM-positive telomere foci. **b,** Quantification of (**a**), showing the number of BLM-positive telomere foci per cell in G2/M-synchronized IMR90-T and IMR90-ALT cells. Data represent mean ± s.e.m. from three independent experiments. Statistical analysis was performed using an unpaired two-tailed Student’s t test; P values are shown. **c,** Western blot analysis of PAGR1, DAXX, STAG2, and PAXIP1 protein expression in parental U87-cdDAXX#1 (Ctrl) cells and two CRISPR-derived PAGR1 deletion clones (dPAGR#1 and #2) transduced with either control (−) or Flpe construct (+). **d,** Telomere dot-blot analysis of anti-STAG2 ChIP in parental U87-cdDAXX#1 (Ctrl) cells and dPAGR clones (#1 and #2) transduced with either control (−) or Flpe construct (+).

